# Spatial patterning of the epigenome during vertebrate gastrulation

**DOI:** 10.1101/2024.02.26.581880

**Authors:** Ana Paula Azambuja, Megan Rothstein, Tatiane Y. Kanno, Marcos Simoes-Costa

## Abstract

A central question in developmental biology is how embryonic cells acquire and store positional information during pattern formation. In vertebrates, this process begins with the localized activation of signaling systems that mediate axial specification. How these spatial cues are recorded within the regulatory landscape of cells has remained unclear. Here, we report that the chromatin landscape of embryonic cells is spatially patterned during gastrulation. Using spatially resolved genomic analysis in avian embryos, we observed that the epigenome becomes organized in gradients of accessibility and activity along the embryonic axes. These chromatin gradients are established within the loci of developmental genes at the onset of gastrulation and can be used to infer the position of cells in the embryo. Our results show that axial specification involves the spatial organization of the epigenome, linking patterns of chromatin activity to the emergence of the embryonic body plan.

## Introduction

Embryonic patterning depends on molecular systems that convey spatial cues to developing cells. This requires the generation of positional information, a set of molecular coordinates that define where cells reside within the embryo^1–3^. Positional information has been a fundamental concept in developmental biology, offering a framework to explain how progenitor cells are patterned in space. It was outlined by Lewis Wolpert^1^, based on his work in Hydra regeneration and on classical experiments performed by Hans Driesch, Thomas Hunt Morgan, Hildegard Stumpf, and others^4^. Wolpert postulated that pattern formation is initiated by the establishment of gradients of signaling molecules along the embryonic axes, known as morphogens^5^. Cells that are closer to the source of the morphogen receive a higher concentration of the molecule, while cells further away receive a lower concentration^6^. These gradients provide spatial context, allowing cells to adopt distinct molecular identities depending on their position^1,3^. This framework anticipated several landmark discoveries, including the identification of the first morphogen in Drosophila^7^ and the demonstration that extracellular signals can direct the activation of gene regulatory networks that determine regional cell identity^8,9^.

Much of the positional information required for the regionalization of the early embryo emerges during axial specification^10^. This refers to the establishment of molecular asymmetries along the anterior-posterior (AP), dorsal-ventral (DV), and medial-lateral (ML) axes in the developing embryo. In *Drosophila melanogaster*, gradients of maternally derived transcription factors establish the embryonic axes, resulting in a system of spatial coordinates across the syncytial embryo^7,11,12^. In vertebrates, which develop from a cellularized blastoderm, axial specification is driven primarily by intercellular signaling. WNTs, BMPs, FGFs, and retinoic acid have all been implicated in establishing embryonic asymmetries^13–15^. Ligands are produced by specific cells and generate concentration gradients via passive or active diffusion, and cells respond to these gradients by activating different sets of genes^8^. Varying concentrations of signals and patterning genes are thus thought to encode positional information^2^. However, recent studies have questioned whether morphogen gradients alone are sufficient to account for the precision and stability of early patterning, suggesting that additional regulatory mechanisms may be involved^16^.

A cellular process that may play a pivotal role in embryonic patterning is the regulation of the chromatin landscape. As early as 1969, long before the characterization of mechanisms of epigenetic regulation, Wolpert proposed that the establishment of positional information would require a coordinate system of spatial genomic responses in the embryo^1^. Yet, despite decades of research, how the epigenome is remodeled during early pattern formation remains poorly understood. A major obstacle has been the technical challenge of profiling chromatin organization in a spatially resolved manner. To address this, we leveraged the large size and accessibility of the avian embryo to perform spatially resolved genomic analyses during gastrulation. This approach enabled us to map chromatin accessibility and gene expression along the embryonic axes with spatial resolution. Our results show that axial specification involves the spatial patterning of the epigenome and that chromatin accessibility changes according to cellular position. We find that a subset of accessible cis-regulatory elements displays regionalized activity that can be used to infer where individual cells reside within the embryo and to reconstruct spatial patterns of gene expression. These findings suggest that early embryonic cells capture positional cues within their chromatin landscape through deployment of specialized cis-regulatory elements.

## Results

### Reconstruction of gene expression patterns in the early avian embryo

To map the transcriptional signatures that underlie axial specification, we conducted a genome-wide reconstruction of gene expression patterns in the early chick embryo. Like its human counterpart, the avian embryo develops as a flat, trilaminar disc, but its larger size makes it particularly well-suited for spatially resolved transcriptomic analysis. We performed Tomography-Seq (tomo-seq)^17^ on embryos at the end of gastrulation (Hamburger-Hamilton stage 5, HH5)^18^, which were cryosectioned along the AP and ML axes (Fig. 1a). Individual 100um tissue sections were then subjected to 3’ RNA-seq (Fig. 1b). Tomo-seq allowed for genome-wide quantification of gene expression levels along embryonic axes, with normalized mRNA levels successfully recapitulating the axial domains of genes like *OTX2*, *MEOX2* and *TBXT* (Fig. 1c). Integration of the AP and ML tomo-seq datasets allowed us to reconstruct gene expression patterns in the avian embryo (Fig. 1d) and to simulate *in situ* hybridization experiments *in silico* for hundreds of genes (Supplementary Fig. 1). Inferred gene expression patterns were consistent with those of known axial-restricted genes (Fig. 1c-e, Supplementary Fig. 2a), including *OTX2*, *CHRD* and *WNT5A*. Identification of genes with similar expression patterns also allowed us to delineate gene signatures for morphological landmarks like the anterior neural plate, primitive streak, and the notochord (Supplementary Data 1).

**Fig. 1.**
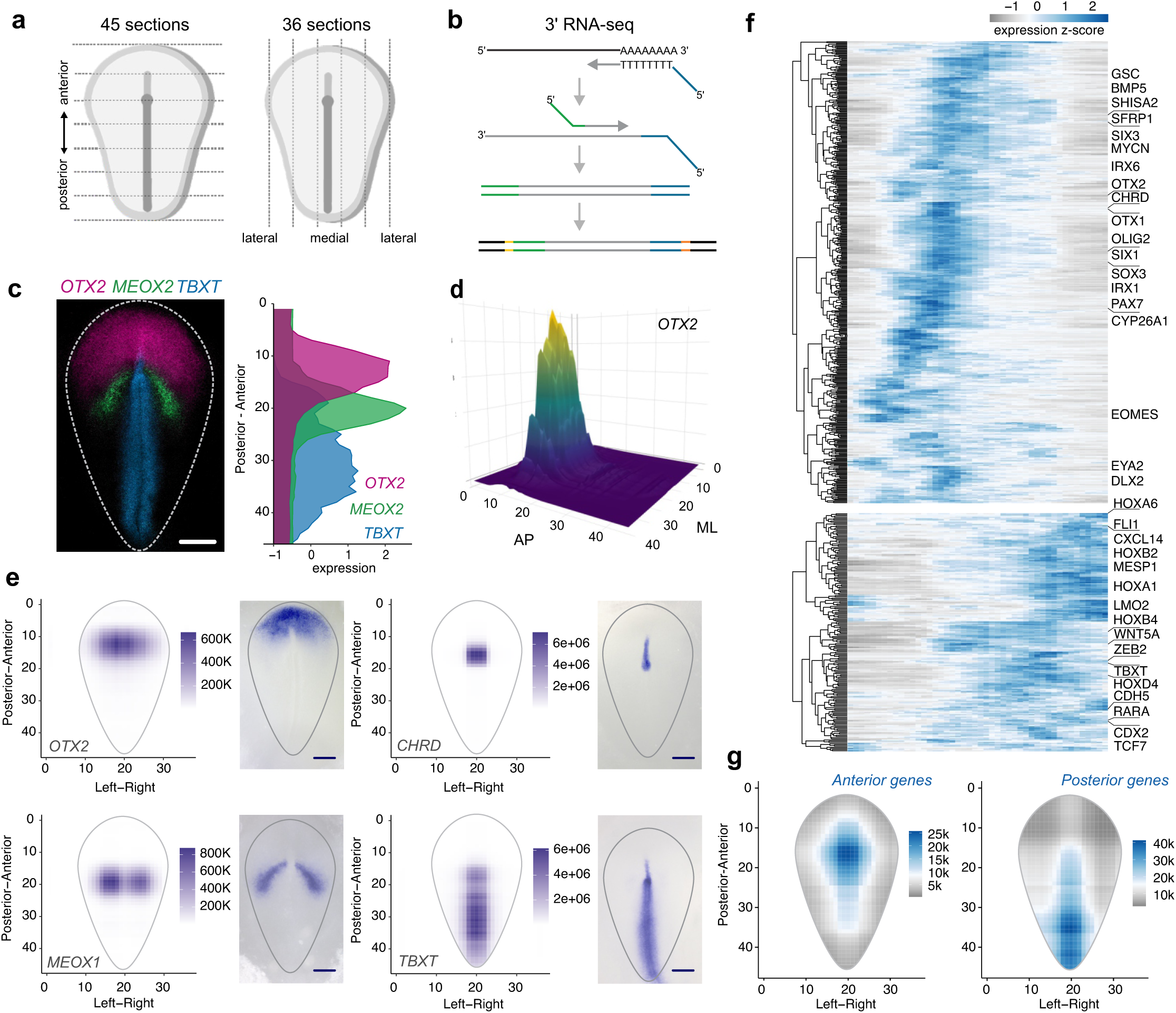
Reconstruction of gene expression patterns in the avian gastrula. (a) Spatial transcriptomic analysis was performed via cryosectioning 100um sections of late gastrula stage (HH5) embryos along the anteroposterior (AP) and mediolateral axes (ML). (b) Individual tissue sections were subsequently processed for 3’ RNA-seq. (c) Multiplexed HCR *in situ* hybridization showing the expression of *OTX2, MEOX2* and *TBXT* in the HH5 chicken gastrula. Density plots to the right display normalized gene expression profiles along the AP axis derived from analysis of the tomo-seq datasets. Scale bar: 400μm. (d) Integration of AP and ML transcriptomic datasets allows for quantification of gene expression in two dimensions across the avian gastrula. The two-dimensional expression of *OTX2* is shown for reference (e). (e) *In silico* (left) reconstruction of the *OTX2*, *CHRD*, *MEOX1*, and *TBXT* 2D gene expression patterns, along with the corresponding *in situ* hybridization images (right). *In silico* projections recapitulate the gene expression profiles as seen via *in situ* hybridization. (f) Heatmap showing 706 anterior-and 294 posterior-enriched genes (p-value <0.01, log fold change >1/<-1) expressed in the avian gastrula, as determined via TradeSeq’s startVsEndTest function. (g) Spatial averages of anterior and posterior genes in the gastrula. Source data are provided as a Source Data file. AP: anteroposterior, ML: mediolateral.

Next, we used the tomo-seq datasets to identify genes with spatially patterned expression along AP axis. Transcriptomes of 100um tissue sections from two biological replicates showed strong concordance between individuals (Supplementary Fig. 2d,e). Hierarchical clustering of gene expression profiles identified distinct clusters of gene expression profiles along the AP axis (Supplementary Fig. 2b). We detected expression of approximately 13,500 genes (>50 counts across all slices), of which 1000 displayed a statistically significant association between gene expression and position along the AP axis. Of these, 706 genes were found to be highly enriched at the anterior region of the embryo (Supplementary Data 2), including previously characterized anterior factors such as *OTX2*, *SIX3* and *CYP26A* (Fig. 1f, Supplementary Data 2)^19–21^. In contrast, only 294 genes were found to be posteriorly enriched. This cohort of genes contained *HOX* family members, and other posteriorizing factors like *CDX4* and *WNT5A* (Fig. 1f, Supplementary Data 2)^22,23^. Gene ontology (GO) analysis of anterior-enriched gene sets showed an enrichment of terms associated with neurogenesis, head development, and sensory structure morphogenesis, while those associated with posterior-enriched genes included mesenchymal tissue formation, including heart and skeletal morphogenesis (Supplementary Fig. 2c). Averaging *in silico* expression patterns of all anterior genes showed that their expression was the strongest around the Hensen’s node, whereas posterior genes were enriched around the primitive streak (Fig. 1g). Together, these results allowed for a high-throughput reconstruction of gene expression patterns along the AP and ML axes and led to the identification of AP gene signatures that are established during avian gastrulation.

### Spatial patterning of the epigenome during axial specification

Next, we examined the organization of the epigenome along the AP axis of the gastrula. We divided HH5 avian embryos into seven transverse sections along the AP axis and subjected them to ATAC-seq analysis^24^ (Fig. 2a). Quality control analysis of our ATAC-seq datasets showed an expected nucleosomal distribution of ATAC-seq fragments, as well as enrichment at transcription start sites (Supplementary Fig. 3a). This tomoATAC-seq strategy led to the identification of a set of cis-regulatory elements (CREs) that display a graded pattern of chromatin accessibility along the AP axis (Fig. 2b). These genomic regions, which we termed regional cis-regulatory elements (rCREs), were particularly abundant in the loci of genes with regionalized expression patterns (Supplementary Fig. 3b). We identified a significant association between genes enriched for rCREs in their loci and those with anterior/posterior enriched expression patterns (*P*=3.635e-06, chi-square test). For instance, the *OTX2* locus contained rCREs that were highly accessible at the anterior-most sections but displayed a gradually weaker ATAC signal in more posterior sections (Fig. 2c). The converse was observed in the locus of *WNT5A*, which contained many posterior rCREs (Fig. 2d). Quantification of normalized chromatin accessibility confirmed that, rather than recapitulating transcriptional domains, rCREs formed stereotypical gradients of accessibility along the AP axis (Fig. 2e,f, Supplementary Fig. 4-5). Chromatin gradients were not uniform within a given locus, as rCREs were often interspersed by elements that were not patterned or that were patterned in the opposing direction along the AP axis (arrows, Fig. 2c,d). Hierarchical clustering of the ATAC-seq datasets revealed varying patterns of chromatin accessibility along the AP axis (Supplementary Fig. 3c). We next compiled anterior and posterior sets of rCREs to assemble AP epigenomic signatures. Differential accessibility analysis identified two sets of 10913 and 12532 elements that were significantly enriched at the anterior and posterior poles of the embryo, respectively, which we deemed anterior and posterior rCREs (Fig. 2b, Supplementary Fig. 3d; Supplementary Data 3). Gene onthology (GO) analysis of the genes associated with anterior or posterior rCREs revealed distinct functional enrichments corresponding to developmental programs operating in each region (Supplementary Fig. 3e). Anterior rCRE-associated genes were enriched for biological processes related to BMP signaling, axis specification, and sensory organ development. In contrast, posterior rCRE-associated genes were linked to processes such as cell motility, embryonic patterning, and tissue morphogenesis.

**Fig. 2.**
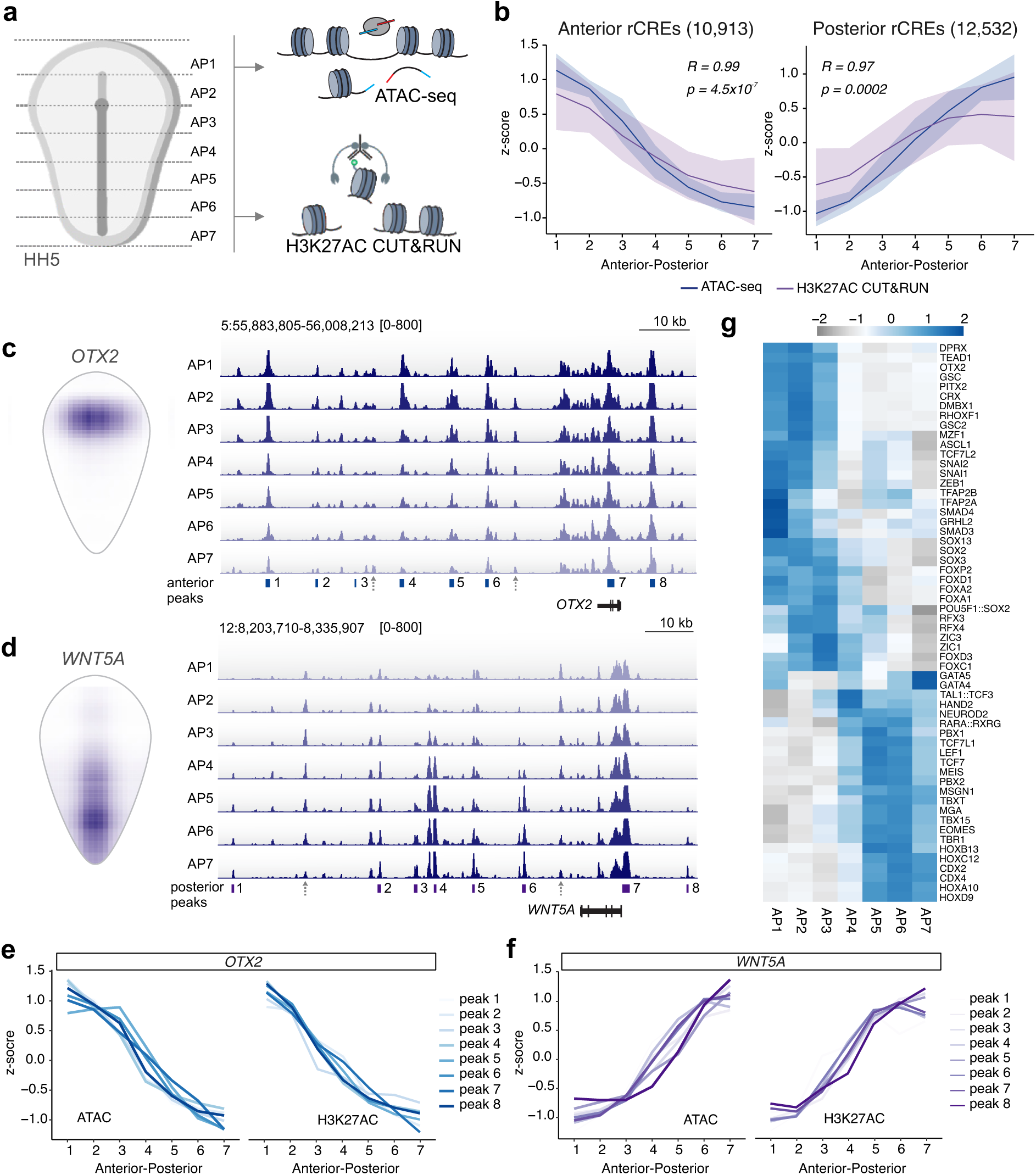
Identification of gradients of chromatin activity in regional cis-regulatory elements (rCREs). (a) using the Hensen’s node as a landmark, HH5 avian embryos were divided into seven sections along the AP axis and subjected to ATAC-seq and CUT&RUN for H3K27AC. (b) Profiles of chromatin accessibility and H3K27AC enrichment of anterior and posterior rCRES (10,913 and 12,532 peaks, respectively) along the AP axis. Lines represent the average ATAC-seq and H3K27AC signal, while ribbons represent standard deviation. The Spearman’s correlation coefficient represents the correlation between the mean ATAC-seq and H3K27AC CUT&RUN values (solid lines). Chromatin accessibility and H3K27AC enrichment are strongly correlated along the AP axis. Correlation assessed using a Pearson Correlation test. (c) *In silico* reconstruction of *OTX2* gene expression alongside ATAC-seq profiles in the *OTX2* locus. Note the gradients of accessibility that are formed from the anterior to the posterior regions of the embryo. Anterior peaks within 100kb of the *OTX2* transcription start site are shown for reference. Arrows highlight regions of chromatin accessibility that are not patterned. (d) *In silico* reconstruction of *WNT5A* gene expression alongside ATAC-seq profiles in the *WNT5A* locus. Posterior peaks within 100kb of the *WNT5A* transcription start site are shown for reference. Accessibility profiles display a posterior to anterior gradient. Arrows highlight elements that are not patterned or with anterior-enriched accessibility. (e) Quantification of rCRE accessibility and H3K27AC enrichment in eight peaks in the *OTX2* locus. Peaks correspond to those shown in panel (c). (f) Quantification of rCRE accessibility and H3K27AC enrichment in eight peaks in the *WNT5A* locus. Peaks correspond to those shown in panel (d). (g) Heatmap displaying variable motifs enriched along the AP axis as identified via Chromvar. Source data are provided as a Source Data file. AP: anteroposterior, HH: Hamburger Hamilton.

To identify the main regulators of AP identity in the embryo, we conducted transcription factor motif enrichment analysis with HOMER^25^ on anterior and posterior rCRE sets. Peaks active at the anterior end of the embryo were enriched for motifs for OTX2, GSC, SOX3, and FOXA2, while those active at the posterior end of the embryo contained HOX, CDX2, and TBXT motifs (Supplementary Fig. 3f). Analysis of transcription factor motif activity across the seven ATAC-seq sections via chromVAR^26^ confirmed high variability of anterior and posterior transcription factor binding motifs along the AP axis (Fig. 2g). Furthermore, we found a strong correlation between chromatin accessibility and deposition of H3K27AC at anterior and posterior rCREs, confirming that these genomic regions are not only open, but also display polarized chromatin activation in the embryo (Fig. 2b, Supplementary Fig. 3g). Similar to chromatin accessibility, H3K27AC deposition in rCREs was also graded along the AP axis (Fig. 2b,e-f, Supplementary Fig. 4-5). Together, these results reveal that the chromatin landscape of the gastrula is spatially patterned along the AP axis, with gradients of accessibility and activity enriched at *loci* of regionally expressed genes. These patterns involve transcription factors known to regulate axial specification and suggest that the epigenome reflects positional cues present in the early embryo.

### Gradients of regional CREs are shaped by signaling pathways

To determine when rCRE gradients first appear during development, we analyzed chromatin patterning during blastula and gastrula stages (Fig. 3a). HH1 (specifically EGK XIII^27^), HH2, and HH3 embryos were divided in five sections along the AP axis and subjected to CUT&RUN for H3K27AC. When visualizing trends of chromatin activity at anterior and posterior rCREs, we observed little to no chromatin polarization at the early (EGK XIII) and late (HH2) blastula stages. Graded activation of rCREs became conspicuous at HH3, coinciding with onset of gastrulation (Fig. 3a, Supplementary Fig. 6a). To verify how rCRE polarization correlates with transcription, we also performed RNA-seq along the AP axis in these early stages. The results showed that expression of the tomo-seq AP gene sets (Fig. 1) became regionalized following the establishment of rCRE gradients (Supplementary Fig. 6a). To confirm that embryos were positioned correctly prior to sectioning, we examined the *NODAL* gene locus. At all three stages assayed, *NODAL* transcripts were found to be enriched in the posterior-most sections (Supplementary Fig. 6b), consistent with its expression in Kohler’s sickle and the early primitive streak^28,29^. To further explore the relationship between early chromatin patterning and transcriptional activation, we examined individual loci of patterning genes that were transcriptionally inactive at the stages of rCRE activation. We observed that in such genes, including *DMBX1* and *HOXA11*, rCRE gradients were established prior to regionalized transcription (Supplementary Fig. 7a,b). These observations indicate that chromatin accessibility gradients can form ahead of gene expression, suggesting that rCRE activation may help prime regulatory landscapes for subsequent transcriptional responses during axis formation. We next examined the mechanisms that shape chromatin gradients during gastrulation.

**Fig. 3.**
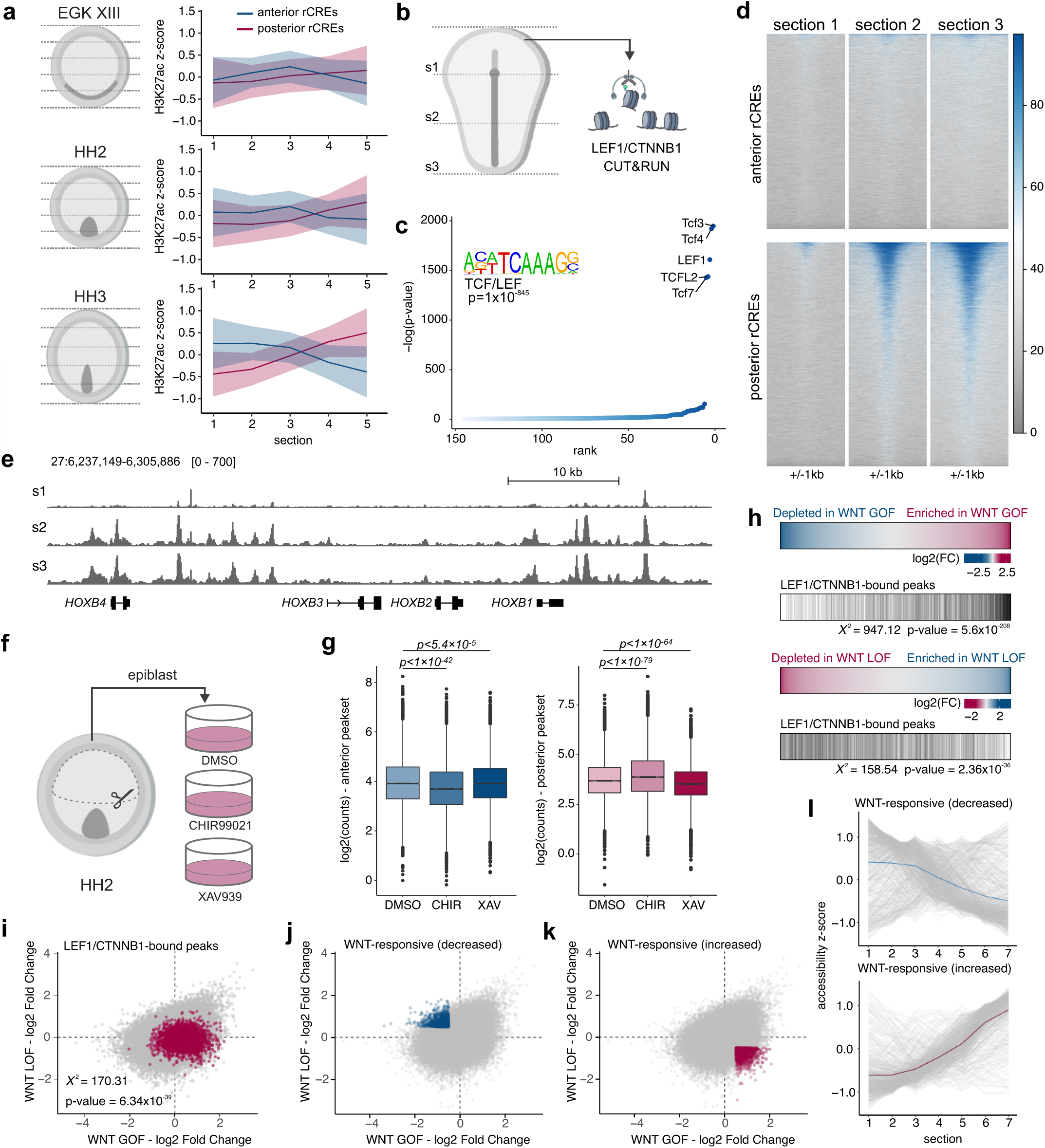
Regulation of rCRE gradients by signaling systems. (a) H3K27AC signal at rCREs along the anterior/posterior axis at three developmental stages. Lines represent the average H3K27AC signal, while ribbons represent standard deviation. Anterior-posterior gradients are formed at the onset of gastrulation (HH3). (b) Strategy for genome-wide mapping of LEF1/CTNNB1 (n=3) across the HH5 gastrula. (c) Motif enrichment analysis of LEF1/CTNNB1 peaks. Top enriched motifs include the WNT nuclear effectors TCF/LEF. (d) Tornado plots displaying LEF1/CTNNB1 signal at anterior-posterior rCREs. Binding of LEF1/CTNNB1 is enriched in posterior rCREs in an anterior-to-posterior gradient. Average signal across three biological replicates is shown. (e) Genome browser tracks showing LEF1/CTNNB1 binding in the *HOXB* cluster. (f) Strategy for manipulation of WNT signaling in naïve avian blastoderm. Cells from anterior epiblast (HH2) were cultured in media supplemented with a WNT agonist (CHIR99021), a WNT antagonist (XAV939), or DMSO as a control. (g) Box plots showing distribution of ATAC-seq signal in anterior/posterior rCREs upon WNT manipulation. Statistical significance assessed using a two-tailed unpaired t-test. Boxplot center lines are median, box limits are upper/lower quartiles, whiskers are the 1.5X interquartile range, and points are outliers (n=3). (h) Heatmaps showing differentially accessible peaks upon WNT GOF (top) and LOF (bottom). The overlap with LEF1/CTNNB1 peaks is indicated by vertical bars. A two-way chi-squared test was used to test the association of LEF1/CTNNB1 peaks and differential accessibility (log2FC>/<0.5). (i) Scatter plot of all accessible peaks across ATAC-seq datasets. Log2 fold change in accessibility upon WNT GOF or LOF is shown on the x and y axes, respectively. LEF1/CTNNB1-associated peaks (highlighted) are significantly associated with WNT-responsive peaks. Statistical significance assessed using a two-way chi-squared test for association between LEF1/CTNNB1 binding and WNT responsiveness. (j-k) Scatter plot as shown in (i), with highly WNT-responsive peaks highlighted. (l) Profiles of chromatin accessibility of WNT-responsive peaks highlighted in (j) (top) and (k) (bottom) from the HH5 tomoATAC-seq dataset. Solid lines represent the mean of all profiles shown. WNT-responsive elements display accessibility gradients along the anterior/posterior axis. Source data are provided as a Source Data file. HH: Hamburger Hamilton, GOF: gain-of-function, LOF: loss-of-function.

A well-known positional cue involved in the establishment of the anteroposterior axis is WNT signaling^14^. This highly conserved signaling system acts as a posteriorizing agent across metazoans^30^. We thus tested the hypothesis that rCREs are directly regulated by nuclear effectors of WNT signaling. Several lines of evidence from the analysis of our tomo-seq datasets point to a role for WNT signaling in the establishment of rCREs. First, a number of WNT ligands and WNT pathway-associated genes are restricted in expression along the AP axis (Supplementary Data 2). Moreover, motif enrichment analysis of posterior rCREs revealed robust enrichment of motifs for nuclear effectors of WNT signaling, including LEF1 and TCF-like factors (Fig. 2g, Supplementary Fig. 3f). To investigate the relationship between rCREs and WNT signaling, we performed CUT&RUN for LEF1/CTNNB1 in three sections along the AP axis of HH5 embryos (Fig. 3b). Analysis of our CUT&RUN datasets revealed 3729 peaks, representing genomic regions bound by LEF1/CTNNB1. Motif enrichment analysis of LEF1/CTNNB1 peaks showed TCF/LEF as the highest ranked motifs, confirming our ability to isolate regions bound by these factors (Fig. 3c). Furthermore, consistent with the posteriorizing role of WNTs, we found that LEF1/CTNNB1 were predominantly associated with posterior rCREs (Fig. 3d), and that CUT&RUN signal displayed an anterior to posterior gradient (Fig. 3d-e). Analysis of earlier embryos (HH3) revealed that the majority of LEF1/CTNNB1 peaks are already present in the early gastrula (Supplementary Fig. 8a,b). Moreover, while we observed greater enrichment of LEF1/CTNNB1 at anterior rCREs at HH3 than HH5, CUT&RUN signal was consistently stronger at the posterior end of the embryo at both stages (Fig. 3d, Supplementary Fig. 8c). Together, these results indicate that posterior rCREs are regulated by components of the WNT signaling pathway.

To test if WNT signaling is consequential for chromatin patterning, we next assessed the effect of modulation of WNT signaling on rCREs. We isolated cells of the anterior epiblast at stage HH2 (prior to rCRE patterning) and cultured the cells for 24 hours under WNT gain-of-function (GOF) and loss-of-function (LOF) conditions (Fig. 3f). The media was supplemented with either a WNT pathway agonist, CHIR99021^31^, a WNT antagonist, XAV939^32^, or DMSO as a control, and cells were subjected to ATAC-seq following treatment. Analysis of anterior- and posterior-enriched rCREs revealed a significant change in accessibility upon WNT modulation. We observed a decrease in the accessibility of anterior CREs upon treatment with CHIR99021, and a concomitant increase in accessibility following XAV939 treatment (Fig. 3g, Supplementary Fig. 8d). The opposite trend was observed for posterior CREs, which displayed an increase in accessibility following CHIR treatment and a decrease in accessibility upon treatment with XAV939. These results demonstrate that posteriorizing signals enhance chromatin accessibility at posterior rCREs while suppressing accessibility at anterior sites.

We next examined how peaks bound by LEF1/CTNN1B responded to manipulations in WNT signaling. We projected LEF1/CTNN1B-bound peaks onto our WNT modulation epigenomic dataset (Fig. 3h,i). As expected, genomic regions associated with LEF1/CTNN1B binding were positively regulated by WNT signaling (i.e., their accessibility is promoted by CHIR99021 and repressed by XAV939). The integration of these datasets allowed us to compile a list of genomic targets of canonical WNT signaling that respond to the manipulation of the pathway (Supplementary Data 4). Finally, we examined the spatial patterns of accessibility in WNT-responsive genomic regions to examine if they display features of rCREs. Projection of the accessibility of WNT-responsive peaks (Fig. 3j,k) in our tomoATAC-seq datasets showed that they display AP polarization and graded accessibility along the AP axis (Fig. 3l). As expected, peaks activated by WNTs formed posterior to anterior gradients, while peaks inhibited by the signaling system are anteriorly polarized (Fig. 3l). These results show that signaling systems modulate AP gradients of chromatin accessibility, linking signaling pathway activity to the spatial organization of the epigenome.

### Chromatin gradients enable inference of cellular position along the embryonic axis

The finding that rCREs are established during gastrulation led us to postulate that these elements capture positional cues in embryonic cells. To test this, we first examined if rCRE signatures (Supplementary Data 3) can be used to infer the position of embryonic cells along the AP axis. We thus performed single cell multiome analysis^33^ of the avian gastrula using the 10X genomics platform to measure both chromatin accessibility and gene expression in individual cells (Fig. 4a). This analysis allowed us to estimate cellular AP position based on rCRE accessibility, and to subsequently test the accuracy of this prediction based on the gene expression signatures from the tomo-seq analysis (Fig. 1). Upon filtering for high quality cells (n=7750) and performing dimensionality reduction, we identified 16 distinct clusters within the multiome dataset (Fig. 4b). Clusters were distributed similarly within the UMAP projection regardless of which dataset was used for dimensionality reduction (RNA-seq, ATAC-seq, or both) (Supplementary Fig. 9).

**Fig. 4.**
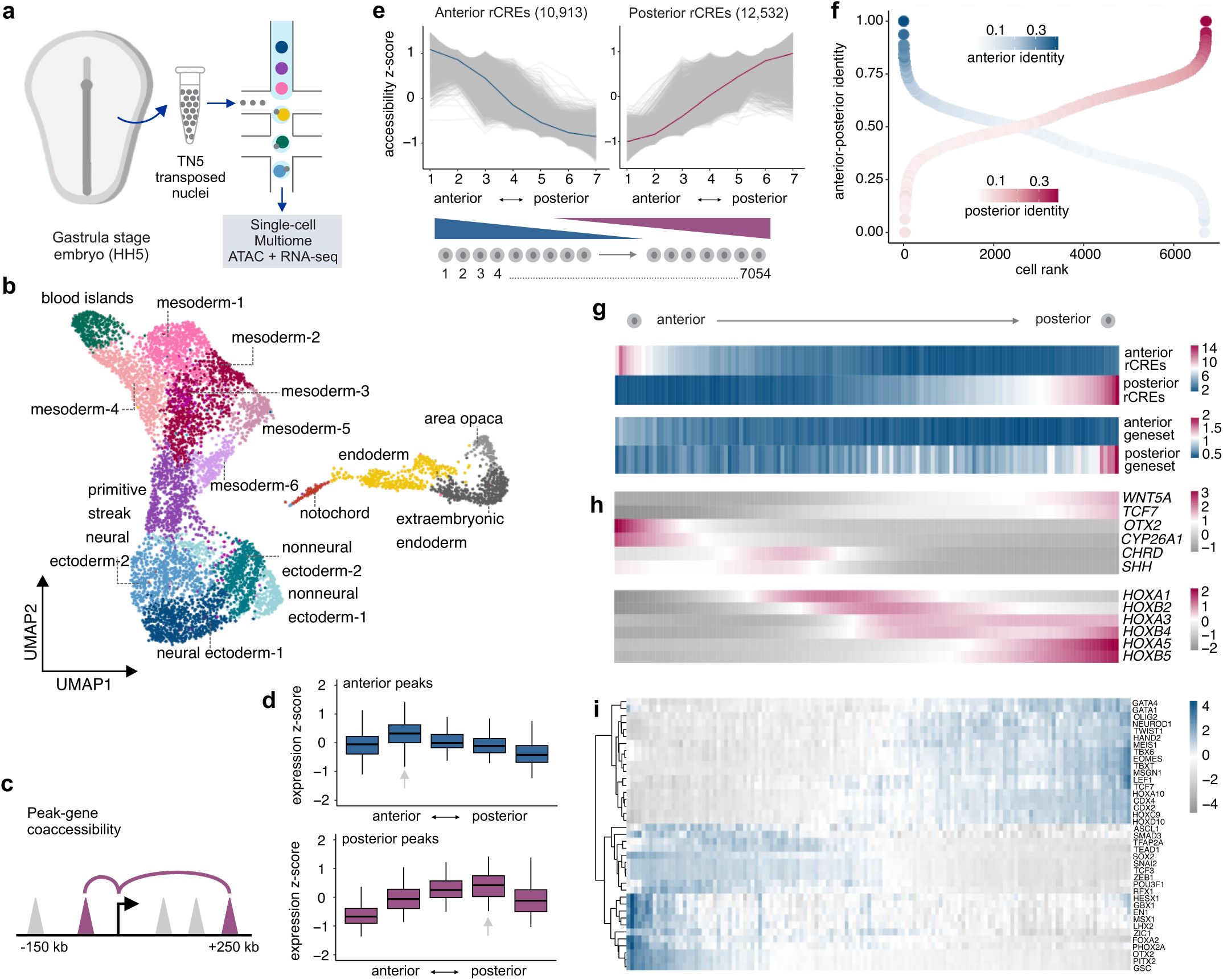
Regional CREs enable inferring the position of cells in the embryo. (a) Single-cell multiomic analysis (RNA-seq & ATAC-seq) of the HH5 chicken embryo via 10X Genomics. (b) UMAP projection of 7750 cells in the avian gastrula, with sixteen distinct clusters labeled by cellular identity. (c) Assignment of rCREs to putative target genes using co-accessibility analysis. (d) Boxplot displaying aggregate AP gene expression profiles via tomo-seq of genes displaying strong co-accessibility with anterior and posterior rCREs. Boxplot center lines are median, box limits are upper and lower quartiles, and whiskers are the 1.5X interquartile range. (e) Strategy to assign cellular position of cells from the multiome dataset based on anterior and posterior rCRE scores. (f) Dotplot displaying fractional identity of individual cells along the AP axis. Cells are colored by similarity to the anterior/posterior rCRE datasets and are ranked from anterior to posterior. (g) Heatmaps displaying accessibility of the anterior and posterior rCRE peaksets (Supplementary Data 3), as well as average expression of anterior/posterior gene sets (Supplementary Data 2) in 50-cell bins ordered along the AP axis. Aggregate scores of anterior and posterior genes are consistent with cell order, showing that rCRE scores can be used to infer cell position. (h) Projection of normalized expression of individual genes from the multiome RNA-seq in 50-cell binds ordered along the AP axis. (i) Chromvar variability of transcription factor motifs in 50-cell binds along the AP axis. Source data are provided as a Source Data file. HH: Hamburger Hamilton.

Cellular identities were assigned to individual clusters using a combination of cluster-specific marker gene expression from scRNA-seq, along with motif enrichment analysis from scATAC-seq (Supplementary Fig. 10A-C). Cluster annotation showed separation of cell identity primarily based on their germ layer of origin. Of the 16 identified clusters, six clusters were mesodermal, four clusters were ectodermal, and one cluster was endodermal (Fig. 4b).

Extraembryonic tissues, including blood islands, extraembryonic endoderm, and the area opaca were also identified in the dataset. After cluster assignment, we attempted to infer the position of individual cells using our anterior and posterior epigenomic signatures. First, we examined whether anterior and posterior rCREs were associated with regionalized gene expression. We employed peak-gene co-accessibility analysis^34^ in the multiome dataset to assign rCREs to their cognate target genes (Fig. 4c) and visualized the expression profiles of these genes along the AP axis. As expected, genes found to be co-accessible with AP rCREs displayed spatial enrichment of anterior or posterior tomo-seq gene signatures (Fig. 4d). Consistent with these findings, GO enrichment analysis of genes linked to anterior and posterior rCREs revealed region-specific developmental signatures (Supplementary Fig. 10D), such as sensory organ and brain development in anterior regions and somitogenesis and mesoderm formation in posterior regions. Next, we attempted to order cells along the AP axis based on epigenomic information.

We utilized a quadratic programming approach^35,36^ to calculate the fractional identity of each cell based on its similarity to the anterior vs. posterior rCRE signatures (Fig. 4e). This strategy allowed us to assign the approximate position of each cell based on rCRE score (Fig. 4f). To test the accuracy of rCRE-based cell ordering, we projected transcriptomic information from the multiome on the ordered cells. We first assessed the expression of the anterior- and posterior-enriched gene sets identified in the tomo-seq analysis (Fig. 1f, Supplementary Data 2). We found that these gene sets were enriched in the anterior and posterior positioned cells, validating our cell ordering analysis (Fig. 4g). Reconstruction of expression patterns of individual AP genes, including *OTX2*, *WNT5A*, and *TCF7*, were consistent with their expression domains (Fig. 4h). This was also true for genes expressed in intermediate AP positions within the embryo, such as *CHRD* and *SHH,* indicating that rCRE gradients contain positional information along the entire embryonic axis. Cell ordering was also able to reconstruct the early *HOX* code in the gastrula (Fig. 4h). Expression patterns were consistent with chromVAR analysis, which identified enrichment of motifs for regulators at their estimated expression domains (Fig. 4i).

Together, these results demonstrate that rCREs can be used to infer the axial position of embryonic cells in silico and support the conclusion that spatial information can be detected in the chromatin landscape.

### Regional CREs provide a spatial register of cells within germ layers

Our multiome analysis demonstrated that the avian gastrula is composed of 16 different cell types (Fig. 4b). We next tested whether we could use the combination of rCRE signatures and quadratic programing to define how these cell types are organized along the AP axis. We first examined the cluster identity of cells enriched for anterior vs. posterior rCREs. We found that the anterior cells are largely ectodermal and endodermal, while the posterior cells consist mostly of mesoderm (Fig. 5a). Indeed, projection of the anterior rCREs on the single-cell multiome UMAP shows robust signal in the ectodermal and endodermal clusters, whereas the posterior signal is strongest in the mesodermal cluster (Fig. 5b). To examine AP composition of the embryo in more detail, we generated 50-cell bins ordered from the anterior to the posterior poles of the embryo. We then assigned the identity of the cells in each bin to determine the regional abundance of distinct cell types (Fig. 5c). This confirmed that a larger percentage of ectodermal and endodermal cells were positioned in the anterior half of the embryo, while mesodermal clusters were posteriorly enriched. To verify the accuracy of this inference, we performed immunostaining in sagittal sections of the avian gastrula with antibodies for markers for the three germ layers (Fig. 5d)^37–39^. Consistent with the cellular composition analysis, endodermal (SOX17+) and ectodermal (SOX2+) progenitors were enriched in the anterior region of the embryo, whereas the posterior gastrula is predominantly mesodermal (TBXT+).

**Fig. 5.**
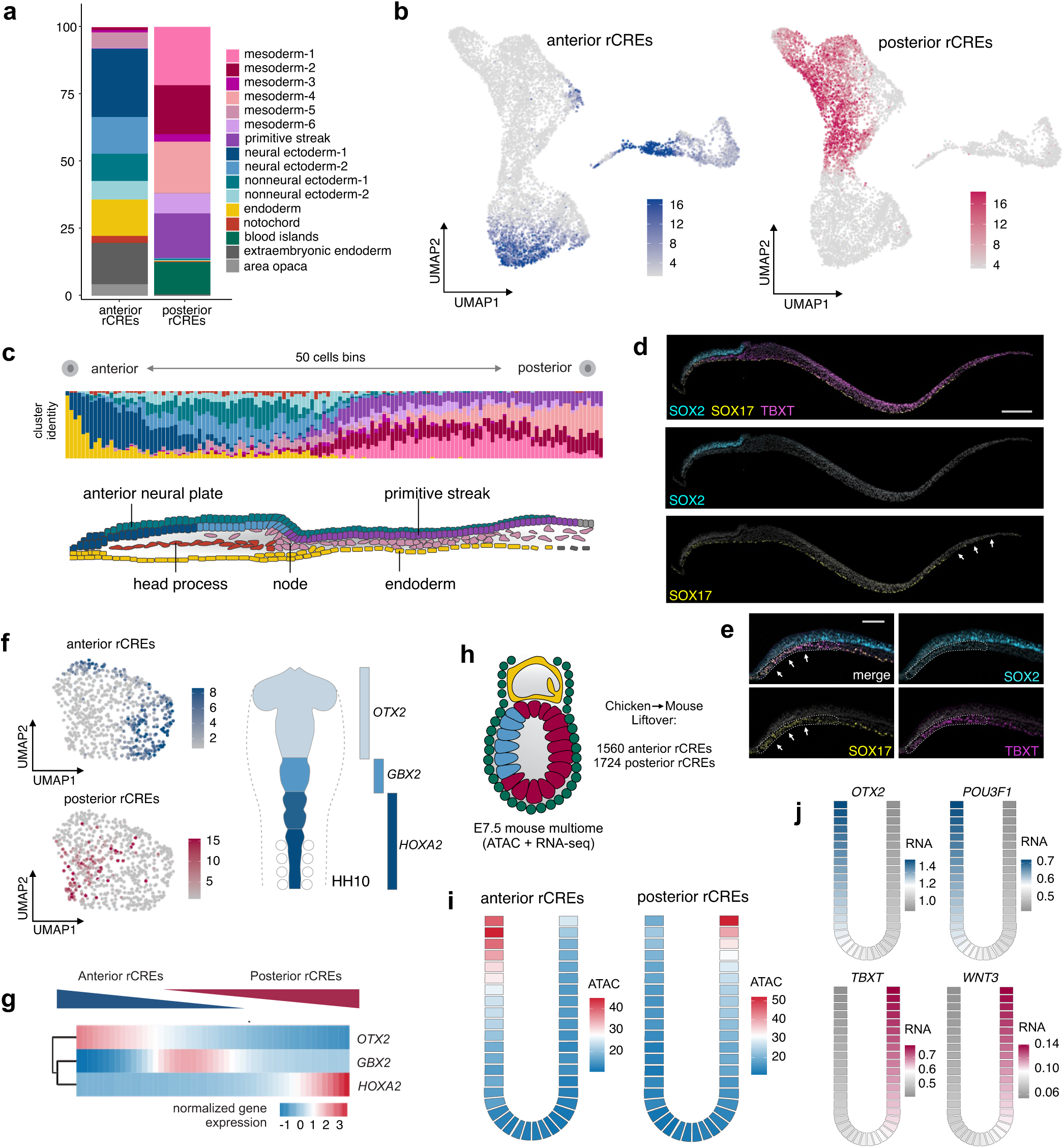
rCREs contain positional information within individual germ layers. (a) Stacked bar plot displaying cellular composition of groups of cells with high anterior vs posterior rCRE scores from the multiome dataset. (b) Projection of rCRE scores in the multiome UMAP. Aggregate accessibility of anterior rCREs is the strongest in ectodermal and endodermal cells, whereas posterior rCREs are on average more accessible in the mesodermal. (c) Cellular composition of 50-cell bins ordered from anterior to posterior using quadratic programming. A schematic sagittal section of the embryo depicts the uneven distribution of germ layers along the AP axis and the position of key midline structures, including the head process. (d) Immunohistochemistry for germ layer markers SOX2, SOX17, and TBXT show ectoderm and endoderm predominantly occupy anterior positions, while most of the posterior embryo is composed of mesoderm (n=3/3). Scale bar: 200μm. (e) Immunohistochemistry for germ layer markers SOX2, SOX17, and TBXT at the anterior end of the embryo (n=3/3). The prechordal plate is outlined and arrows indicate enriched SOX17 expression in this tissue. Scale bar: 100μm. (f) Projection of anterior and posterior rCRE signatures in 708 neuroectodermal cells isolated from the multiome dataset. Axial markers of the neural ectoderm include *OTX2*, *GBX2,* and *HOXA2*. (g) Projection of normalized expression of *OTX2*, *GBX2,* and *HOXA2* in neuroectodermal cells ordered from anterior to posterior using quadratic programming. Cell ordering recapitulates the expected expression patterns for axial markers of the central nervous system. (h) 1560 anterior rCREs and 1724 posterior rCREs are conserved between chicken and mouse. Conserved rCREs were used to order cells from the E7.5 mouse multiome dataset from Argelaguet et al.^43^ as 200-cell bins from anterior to posterior. (i) Heatmaps displaying accessibility of conserved rCREs in 200-cell bins ordered from anterior to posterior in the mouse gastrula. (j) Heatmaps displaying expression of *OTX2*, *POU3F1*, *TBXT*, and *WNT3* in 200-cell bins ordered from anterior to posterior using rCRE accessibility. Conserved rCREs can be used for cell ordering and inference of expression patterns of axial genes in the cup-shaped murine gastrula.

Closer examination revealed that the anterior accumulation of SOX17+ endoderm reflects the presence of the prechordal plate, where the endoderm forms a multi-layered structure. In contrast, the posterior region lacks definitive endodermal cells at this stage, with the underlying layer being largely composed of hypoblast cells (Fig 5e). Furthermore, we also observed that cells assigned to distinct embryonic structures, including the primitive streak and the notochord, occupied the expected position along the AP axis (Fig. 5c).

The asymmetric distribution of cell types along the AP axis led us to test if rCREs could be used to infer the spatial distribution of cells within individual germ layers. We subdivided our multiome dataset into ectodermal, mesodermal, and endodermal clusters (2481, 2936, and 440 cells, respectively) (Supplementary Fig. 11a-c). Projection of the anterior- and posterior-enriched rCREs onto germ layer-specific subclusters revealed distinct subpopulations of cells enriched for axial-restricted peaks in each of the three germ layers (Supplementary Fig. 11a-c). Thus, although unsupervised clustering largely groups cells based on cellular origin, a closer look at intra-cluster organization suggests additional separation based on positional identity. To test this, we examined the axial identity of cells from neural ectoderm, which undergoes early regionalization along the AP axis during the formation of the central nervous system^40^. We employed rCRE-based cell ordering in the neural ectoderm subcluster (Fig. 5f, 708 cells), and assessed the expression of the forebrain, midbrain, and hindbrain markers (*OTX2*, *GBX2*, and *HOXA2*, respectively) (Fig. 5f)^41,42^. Ordered cells displayed the expected expression profiles of these marker genes along the AP axis (Fig. 5g). Next, to determine the extent to which axial-restricted genes are shared across cell types, we conducted trajectory-based differential expression analysis in the ectodermal and mesodermal subclusters (2481 cells and 2936 cells, respectively). This analysis revealed a set of 64 genes significantly associated with AP-restricted expression in both germ layers (Supplementary Fig. 11d,e). Together, these results support the existence of a regulatory code associated with AP patterning that is independent of cellular identity.

### Conservation of rCRE signatures across vertebrates

Spatial epigenomic analysis of the avian gastrula allowed for the identification of anterior and posterior rCREs, which provide a molecular signature of cellular position in the embryo. To determine if this mechanism is present on other vertebrate embryos, we analyzed multiomic data from the E7.5 mouse embryos^43^. Liftover^44^ of anterior and posterior rCREs identified 1560 anterior and 1724 posterior rCREs conserved in mouse (Fig 5h). We then used quadratic programming to assign the relative position of individual cells from the E7.5 mouse dataset along the AP axis (Fig. 5i). Upon ordering cells based on their chromatin accessibility at conserved rCREs, we assessed the expression patterns of anterior- and posterior-restricted genes as 200-cell bins along the AP axis. This analysis revealed spatial restriction of anterior genes *OTX2* and *POU3F1*, as well as posterior genes *TBXT* and *WNT3* in the developing mouse gastrula (Fig. 5j)^45^. Therefore, even a small subset (∼15%) of rCREs can be used to estimate anterior/posterior gene expression patterns across species. These findings suggest that regional chromatin organization associated with positional cues is conserved among amniotes and may represent a general strategy for spatial patterning during early development.

## Discussion

For decades, gradients of secreted morphogens and transcription factors have provided the prevailing explanation for how positional information is generated and processed in the early embryo^4^. Our findings expand this framework by showing that the chromatin landscape of embryonic cells is spatially organized. We observed that during gastrulation, regional gradients of chromatin accessibility emerge along the embryonic axes. These patterns encompass a large fraction of accessible chromatin regions and are interspersed with non-patterned cis-regulatory elements, indicating that spatial cues are recorded only in specialized genomic domains. Many of these patterned regions coincide with developmental genes that display restricted expression domains, including areas of dense regulatory activity such as the *HOX* clusters. Chromatin patterning was also evident within individual germ layers, suggesting that distinct cell types operate within a shared spatial framework of genomic regulation. Understanding how these chromatin states interact to generate three-dimensional gene expression patterns will require elucidating how multiple cis-regulatory elements coordinate transcription and how thresholds of accessibility are established and interpreted in specific embryonic contexts.

Extensive cell movements during gastrulation, including polonaise movements^46,47^ and axial elongation^48^, result in the continuous repositioning of cells along the AP axis. These large-scale rearrangements complicate how spatial cues are maintained and interpreted during cell migration. Our data show that rCREs are dynamically regulated by nuclear effectors of key signaling pathways, providing a mechanism for chromatin accessibility patterns to adapt as cells move through changing spatial environments. Specifically, we show that canonical WNT signaling regulates chromatin accessibility in posterior regions, with nuclear effectors binding regulatory elements in graded fashion along the AP axis. Together with the enrichment of motifs for other nuclear effectors, including those of FGF, retinoic acid, and TGFβ pathways^49–51^, these findings indicate that spatial chromatin patterning arises from the integrated activity of multiple signaling systems. Indeed, the rCRE patterns described here do not represent the final positions of embryonic cells, but rather an early regulatory framework that is subsequently refined as morphogenesis progresses. In this view, the epigenome acts as a dynamic readout of positional cues, integrating multiple morphogen inputs as cells experience changing signaling environments.

Our results also provide insight into how anterior and posterior cell identities are established during gastrulation. Classical models of vertebrate axial specification, largely based on neural induction studies, proposed that the nervous system initially forms with an anterior “default” identity that is later posteriorized by trunk signals^52–55^. In contrast, our data show that distinct anterior and posterior chromatin states emerge simultaneously at the onset of gastrulation. These findings are consistent with Otto Mangold’s organizer graft experiments, which demonstrated that separate organizer regions can induce neural tissues with different axial identities^56^, and with more recent work showing that posterior identity can arise before neural induction^53^. Together, these observations suggest that anterior and posterior territories are specified very early in development by the combined action of signaling pathways. The regulation of rCREs by signaling pathways that define these territories provides a molecular explanation for this early patterning, showing how the epigenome connects morphogen activity to region-specific gene expression programs during gastrulation.

## Methods

### Ethics statement

This work was conducted with ethical regulations and guidelines approved by Boston Children’s Hospital (IBC-RN00001327-1).

### Tissue collection

Fertilized Leghorn White chicken eggs were obtained from the Department of Animal Science, University of Connecticut. Eggs were incubated at 37° C and 50% humidity until embryos reached the desired developmental stage. Embryos were collected in Ringer’s solution, staged according to the Eyal-Giladi and Kochav^27^ or Hamburger and Hamilton^18^ staging systems, and prepared for genomic protocols as described below. EGK XIII embryos were collected before primitive streak formation and staged by assessing epiblast expansion and Koller’s sickle position. Embryos were examined and dissected under a ZEISS Stemi 508 stereo microscope. The anterior-posterior and lateral axes were measured, and embryos of similar size were grouped for processing.

### Tomo-seq sample and library preparation

HH5 embryos were collected, staged, and cleaned in RNase-free solution. Similar sized embryos were then transferred to freezing molds, positioned depending on the axis to be sectioned and embedded in frozen section medium (Thermo Scientific™ Richard-Allan Scientific Neg 50™, #6502). Before freezing, each embryo had its Hensen’s node, anterior-posterior and lateral ends marked with dye to serve as landmarks for the sectioning process. In a RNase-free environment, frozen samples were trimmed to avoid extra-embryonic tissues and frozen medium excess, and cryosectioned into 100um sections. Each anterior-posterior or medial-lateral section was then collected for total RNA extraction with the RNAqueous-Micro Total RNA Isolation Kit (Thermo Fisher Scientific, # AM1931) according to the kit’s protocol and processed for 3’ RNA-seq library preparation. Libraries were prepared using the QuantSeq 3’mRNA-seq Library Prep Kit (FWD) for Illumina (Lexogen, #K01596), following the manufacturer’s recommendations for low RNA input samples. Final libraries were quantified, pooled, and sequenced with single-end 75bp reads on an Illumina NextSeq500 instrument.

### Tomo-seq data analysis

Trimming of adapter sequences and poly(A) tails was performed using BBDuk (version 38.93) using the following parameters: ref=polyA.fa.gz,truseq_rna.fa.gz k=13 ktrim=r useshortkmers=t mink=5 qtrim=r trimq=10 minlength=20. Next, STAR (version 2.7.3a)^57^ was used to align trimmed reads to the galGal6 chicken reference genome using the following arguments: --outFilterType BySJout --outFilterMultimapNmax 20 --alignSJoverhangMin 8 --alignSJDBoverhangMin 1 --outFilterMismatchNmax 999 --outFilterMismatchNoverLmax 0.1 -- alignIntronMin 20 --alignIntronMax 1000000 --alignMatesGapMax 1000000 --outSAMattributes NH HI NM MD --outSAMtype BAM SortedByCoordinate. Bam alignment files were indexed using samtools (version 1.10)^58^ and reads were counted with htseq-count (version 0.13.5)^59^ using a GTF file containing chicken gene annotations and using the following parameters: -m intersection-nonempty -s yes -f bam -r pos. AP section 14 from replicate 1 was omitted from downstream analyses because it received very few counts and was deemed a failed library.

Differential gene expression analysis was performed on two embryo replicates using TradeSeq (version 1.8)^60^. Anterior- and posterior-enriched genes were identified using TradeSeq’s startVsEndTest function and genes with a p-value <0.01 and a log fold change >1 or <-1 were considered significantly enriched. For plotting of AP and 2D gene expression profiles, raw counts were normalized to the total number of spike-in reads and the moving average of the normalized counts were plotted along each axis. Gene ontology analysis was performed using PANTHER’s^61,62^ Fisher’s Exact test with Bonferroni correction.

### *In situ* hybridization

For *in situ* hybridization, embryos were fixed in phosphate buffer saline (PBS) containing 4% paraformaldehyde (PFA) for 2 hours at room temperature or overnight at 4 °C. Following fixation, embryos were dissected, washed with PBS containing 0.1% Tween (PBST), dehydrated and stored in methanol at −20° C. Whole-mount in situ hybridization was performed as previously described^63^. Embryos were imaged using a ZEISS Stemi 508 stereo microscope.

### HCR RNA-FISH

Hybridization chain reaction RNA fluorescent in situ hybridization (Molecular Instruments HCR v3.0) was performed in HH5 (19-22h) whole mount chick embryos^64^. Briefly, embryos were collected and fixed in 4% paraformaldehyde for 2 hours at room temperature. Embryos were dissected to remove the vitelline membrane, washed with PBST on ice and then stored in methanol at −20°C until use. Next, embryos were rehydrated with a series of graded Methanol/PBST washes on ice. For the detection step, embryos were pre-hybridized with probe hybridization buffer for 30 minutes at 37°C, then incubated with 4 pmol of each probe set overnight. The excess of probe was removed by washing the embryos with probe wash buffer at 37°C. The embryos were then pre-amplified with amplification buffer and incubated with hairpin solution overnight in the dark at room temperature. Excess hairpins were removed by washing the embryos with 5X SSCT at room temperature. Embryos were incubated with DAPI for 1 hour followed by sample mounting for microscopy. Embryos were imaged at the Nikon Imaging Center at Harvard Medical School using a Nikon AX R point scanning confocal. The target mRNA probe sets, amplifiers and buffers were purchased from Molecular Instruments (https://www.molecularinstruments.com/).

### Immunohistochemistry

For whole-mount immunohistochemistry, embryos were collected and fixed in 4% PFA-PB for 20 minutes at room temperature. Post fixation, embryos were dissected from the filter paper and washed in TBS containing 0.1% Triton and 1% DMSO (TBTD). Embryos were blocked at room temperature for 2 hours in TBTD supplemented with 10% donkey serum and incubated in anti-SOX2 (Abcam, #ab97959) and anti-SOX17 (R&D, #AF1924) primary antibodies diluted in blocking solution, overnight at 4° C. Following the primary antibody incubation, embryos were washed, blocked for 30 minutes at room temperature, and stained with appropriate secondary antibodies and anti-Brachyury Alexa Fluor 488 Conjugated (R&D, #IC2085G) for 2 hours at room temperature. Secondary antibodies used included donkey anti-rabbit/goat IgG conjugated with Alexa Fluor594/647 (Molecular Probes). Following the secondary antibody step, the embryos were washed, stained with DAPI and post-fixed with 4% PFA for 1 hour, prior to imaging. Whole-mount images were collected using an upright ZEISS Axio Imager fluorescent microscope.

### TomoATAC-seq

HH5 chicken embryos were collected in Ringer’s solution and stage- and size-matched embryos were dissected out of any extra embryonic tissue. Then, using the Hensen’s node as a landmark, seven sections were obtained along the antero-posterior axis, three anterior to the node and four posterior to it. For EGK XIII, HH2, and HH3 embryos, staging was based on established morphological criteria, and embryos were similarly matched for size and developmental progression. These earlier embryos were also cryosectioned into five evenly spaced sections along the embryonic axis. Five position-matching sections were pooled and dissociated in Accumax (Innovative Cell Technologies, #AM105). Cell suspensions were washed and processed using the OMNI-ATAC-Seq protocol^65^. Briefly, dissociated cells were washed in resuspension buffer containing 10 mM Tris-HCl pH 7.4, 10 mM NaCl, and 3 mM MgCl2. After washing, cells were resuspended in lysis buffer composed of the resuspension buffer plus 0.1% NP-40 (Sigma, I8896), 0.1% Tween-20 (Sigma, P1379), and 0.01% Digitonin (Millipore, 300410). The samples were incubated on ice for 3 min, followed by washing with resuspension buffer containing only 0.1% Tween-20. Nuclei were then resuspended in 50 ul of transposition mixture containing 10 mM Tris-HCl pH 7.4, 5 mM MgCl2, 10% Dimethyl Formamide (Sigma, D4551), 1X PBS, 0.1% Tween-20, 0.01% Digitonin, and 5 ul Tn5 transposase (Illumina, 20034210). Transposition was performed for 1 h at 37°C in a thermal mixer. The pre-amplification DNA was purified using the Qiagen Minelute Reaction Cleanup Kit (Qiagen, 28204). Library amplification PCR was performed using the Q5 High-Fidelity 2X Master Mix (NEB, M0492S). Library size selection and cleanup was carried out using Ampure XP beads (Beckman Coulter,A63881). Final libraries were quantified, pooled, and sequenced with paired-end 37bp reads on an Illumina NextSeq500 instrument.

### TomoATAC-seq data analysis

Trimming of Nextera adapter sequences was performed using cutadapt (version 3.4)^66^ using the following parameters: -a CTGTCTCTTATACACATCT -A AGATGTGTATAAGAGACAG --minimum-length=25. Trimmed reads were aligned to the galGal6 chicken reference genome using bowtie2 (version 2.4.4)^67^ with the following arguments: --local --very-sensitive-local --no-unal --no-mixed --no-discordant -X 2000. Bam alignment files were indexed using samtools (version 1.10), followed by marking of duplicate reads using Picard’s (version 2.26.2) MarkDuplicates function. Duplicate reads were then removed using samtools view with the following arguments: -F 1804 -f 2. Bigwigs were generated from Bam files using DeepTools’ bamCoverage function (version 3.5.1)^68^ with the following arguments: --binSize 5 --normalizeUsing RPKM --extendReads. All downstream analyses were conducted using peaks called using cluster-specific peak calling on the HH5 multiome dataset (316,412 peaks, see *10X multiome data analysis*). By calling peaks on individual cell clusters, this increased the sensitivity of peak calling and ensured rare, cell type-specific regulatory elements were not omitted from the analysis. TomoATAC-seq reads in these 250,772 peaks were counted using DiffBind (version 3.4.11)^69^. Peaks with anterior- and posterior-enriched accessibility were identified using TradeSeq’s (version 1.16.0) startVsEndTest function and peaks with a p-value <0.01 and a log fold change >1 or <-1 were considered significantly enriched. For plotting of accessibility profiles, raw counts were library size normalized and the moving average of the normalized counts were plotted along the AP axis. Gene enrichment analysis was performed by assigning each peak to the nearest annotated transcription start site using GenomicRange’s^70^ (version 1.54.1) nearest() function. Next, a binomial test was conducted using the total number of *cis*-regulatory elements assigned to a gene as the number of trials and the proportion of all rCREs over the total number of *cis*-regulatory elements as the probability of success. Genes returning a p-value of </=0.05 were considered significantly enriched for rCREs. Motif enrichment analysis was performed by calculating motif deviations using ChromVar (version 1.16)^26^, as well as using HOMER’s^25^ findMotifsGenome.pl function. Functional enrichment analysis of anterior and posterior rCREs was conducted using GREAT^71^ (version 2.4.0).

### Anterior epiblast ATAC-seq

As described previously, chick embryos were obtained from fertile eggs, collected, and staged in Ringer’s solution. HH2 embryos were dissected off any extra-embryonic tissue and had the anterior hypoblast removed. Then, the anterior, naive, epiblasts from three embryos were microdissected, pooled, and transferred to 96-well plates previously coated with fibronectin and containing 100 uL of 10% FBS DMEM culture media containing the WNT agonist CHIR99021 (Tocris, #4423), or antagonist XAV939 (Sigma, #X3004-5MG) at the final concentrations of 10 uM (the same volume of vehicle, DMSO, was used for control explants). The explants were incubated for 24 hours at 37° C in a CO2 incubator. After this period, explants were washed in PBS and collected with Accumax. Cell suspensions were washed and processed using the OMNI-ATAC-Seq protocol^65^ as described above.

### Epiblast ATAC-seq data analysis

Initial processing of ATAC-seq libraries, including trimming, alignment, removal of duplicate reads, peak calling, and generation of bigwigs, was performed in the same manner as the tomoATAC-seq analysis. After initial processing, ATAC-seq reads were counted at 250,772 peaks obtained from the analysis of the 10X multiome data DiffBind (version 3.4.11). Pairwise differential accessibility analysis between the DMSO vs CHIR99021 and DMSO vs XAV939 conditions was performed via DESeq2^72^ (version 1.42.1) using three biological replicates. Wnt-responsive (increased) peaks were defined as those enriched in the CHIR99021 condition (log2 fold change > 0.5) and depleted in the XAV939 condition (log2 fold change < −0.5). Wnt-responsive (decreased) peaks were defined as those enriched in the XAV939 condition (log2 fold change > 0.5) and depleted in the CHIR99021 condition (log2 fold change < −0.5).

### CUT&RUN

As described above for the TomoATAC-seq experiments, the embryos processed for CUT&RUN were first collected and stage- and size-matched. For each experiment, position matched sections from 3 embryos were pooled together. H3K27AC CUT&RUN experiments were performed with EGK XIII, HH2, HH3, and HH5 embryos sectioned into 5, 5, 6 and 7 sections, respectively. Embryonic structures like Hensen’s node and primitive streak were used as landmarks. The sectioning was performed in a way to preserve the amount of tissue per fragment. For the CTNNB1/LEF1 CUT&RUN, embryos from three biological replicates were dissected into 3 anterior-posterior sections. The rostral segment was cut right after the regressing Hensen’s node and the remaining embryo was cut in half, dividing the middle and caudal segments. CUT&RUN experiments were conducted as previously described^73^. First, the tissue was dissociated into a single cell suspension using Accumax at room temperature for 20 minutes under mild agitation. Next, cells were bound to BioMag Plus Concavalin A magnetic beads (Bangs Laboratories, BP531) and incubated with rabbit anti H3K27AC (AbCam, #ab177178), or rabbit anti-CTNNB1 (Abcam, #ab32572) plus anti-LEF1 (Abcam, #ab137872) antibodies overnight at 4°C, rocking. All antibodies were used at a 1:50 concentration. After washing away the unbound antibody, protein A-MNase was added to a final concentration of 700 ng/mL and incubated on the tube rotator for 1 hour at 4°C. Cells were chilled down to 0°C and CaCl2 was added to a final concentration of 2 mM to activate the MNase enzyme. MNase digestion was conducted for 45 minutes and terminated by adding the 2XSTOP buffer containing heterologous *Saccharomyces cerevisiae* spike-in DNA at a concentration of 2 pg/mL. The protein-DNA complexes were released by centrifugation and digested with proteinase K for 10 minutes at 70°C. DNA fragments were extracted using phenol-chloroform and ethanol precipitation. CUT&RUN libraries were prepared using the NEBNext Ultra II DNA Library Prep Kit (New England Biolabs, #E7645) following the manufacturers protocol. Fragment analysis was performed with Agilent 4200 TapeStation to perform quality control for the libraries. Equimolar concentrations of the libraries were pooled and sequenced with paired-end 37bp reads on an Illumina NextSeq500 instrument.

### CUT&RUN data analysis

Trimming of TruSeq adapter sequences was performed using cutadapt (version 3.4) using the following parameters: -a AGATCGGAAGAGCACACGTCTGAACTCCAGTCA -A AGATCGGAAGAGCGTCGTGTAGGGAAAGAGTGT --minimum-length=25. Trimmed reads were aligned to the galGal6 chicken reference genome using bowtie2 (version 2.4.4) with the following arguments: --local --very-sensitive-local --no-unal --no-mixed --no-discordant -I 10 -X 1000. Bam alignment files were indexed using samtools (version 1.10), followed by marking of duplicate reads using Picard’s (version 2.26.2) MarkDuplicates function. Duplicate reads were then removed using samtools view with the following arguments: -F 1804 -f 2. Peaks were called with macs2 (version 2.2.7.1) using the parameters: -f BAMPE -g 1e9 -q 0.05 --call-summits. Bigwigs were generated from Bam files using DeepTools’ bamCoverage function (version 3.5.1) with the following arguments: --binSize 5 --normalizeUsing RPKM --extendReads. Reads in peaks were counted using DiffBind (version 3.4.11). For plotting of chromatin activity profiles, raw counts were library size normalized and the moving average of the normalized counts were plotted along the AP axis. Motif enrichment analysis and identification of the top enriched transcription factor binding motifs was performed using HOMER’s findMotifsGenome.pl function. For LEF1/CTNNB1 CUT&RUN analysis, consensus peaksets from macs2-called peaks were obtained using DiffBind’s dba.count() function with minOverlap=3. Average bigwigs were generated from three replicates using DeepTools’ (version 3.5.1) bigwigAverage function. Plotting of LEF1/CTNNB1 binding scores at anterior-/posterior-enriched peaks was accomplished using DeepTools’ computeMatrix and plotHeatmap functions, using averaged bigwigs as input. LEF1/CTNNB1 peaks were mapped onto ATAC-seq peaks using GenomicRanges findOverlaps() function.

### Tissue isolation and 10X multiome library preparation

The nuclei of three whole chick embryos at stage HH5 were isolated following the Chromium Next GEM Single Cell Multiome ATAC + Gene Expression (GEX) protocol. Briefly, the embryos were dissected and dissociated using Accumax for 15-20 minutes at room temperature to generate a single cell suspension. Next, the cell suspension was filtered using a Flowmi cell strainer (Millipore-Sigma), porosity 40uM to remove cell debris and clumps. To isolate the nuclei, cells were centrifuged at 500 x g for 5 minutes at 4°C and then lysed using a 0.1X lysis buffer for 5 minutes on ice. The isolated nuclei were washed and resuspended in Diluted nuclei buffer. Finally, the nuclei concentration was determined using an automated cell counter and 15,000 nuclei were used to target a 10,000 nuclei recovery. The ATAC and gene expression library preparation and sequencing were performed at Cornell University BRC Genomics Core Facility according to the manufacturer’s instructions (10X genomics). Final libraries were sequenced on an Illumina NextSeq500 instrument.

### 10X multiome data analysis

Initial processing and alignment of paired scRNA-seq and scATAC-seq datasets from the 10X multiome kit was performed using the cellranger-arc pipeline (version 2.0.0)^74^. Upon generation of the UMI count matrix via cellranger-arc’s count function, downstream processing was performed using Seurat (version 4.3.0)^75^ and Signac (version 1.8.0)^34^. In addition to cellranger’s initial cell quality filtering, cells were additionally filtered in Seurat using the following criteria: nCount_ATAC<8e4 & nCount_ATAC>4e3 & nCount_RNA<5000 & nCount_RNA>300 & percent.mt<10. In total, 7750 cells passed cell quality filtering. RNA- and ATAC-seq datasets underwent independent preprocessing and dimensionality reduction using the standard Seurat and Signac pipelines, respectively. Upon initial dimensionality reduction and clustering of the ATAC-seq dataset, peaks were re-called on individual clusters using MACS3^76^ resulting in a final set of 316,412 peaks. The two datasets were then integrated by calculating a weighted-nearest neighbor (WNN) graph, which was used for final UMAP visualization and clustering.

Clusters were empirically assigned using marker genes identified through gene expression and motif enrichment across UMAP clusters. Ordering of cells along the AP axis was accomplished using a quadratic programming approach developed by Yu and colleagues^35^. Extraembryonic clusters were excluded from the cell ordering analysis, as they were not predicted to be restricted along the AP axis. Cells were ranked according to their accessibility at anterior-/posterior-enriched peaks derived from TradeSeq analysis of the bulk tomoATAC-seq datasets. Cell ranks were assigned using custom R scripts from Yu and colleagues^35^. Smoothened gene expression profiles along the AP axis were computed using TradeSeq’s predictSmooth() function. Peak-gene linkage analysis was performed by computing the correlation between gene expression and accessibility at nearby peaks using Signac’s LinkPeaks function. Only high-confidence peak-gene links (p-value < 1×10^-6^) were used for downstream analysis. Analysis of individual germ layers was performed by subsetting the Seurat object to include only the clusters of interest, followed by repeating dimensionality reduction and WNN integration on each germ layer-specific data subset. AP differentially expressed genes in individual germ layers were identified using TradeSeq’s associationTest() function and genes with an fdr <0.05 were considered significantly enriched.

### Mouse multiome data analysis

For the analysis of conserved rCREs in the mouse gastrula, 10913 anterior and 12532 posterior rCREs were lifted over to the mouse mm10 genome using UCSC genome browser’s LiftOver^44^ tool. This resulted in 1560 and 1724 conserved anterior and posterior rCREs, respectively. E7.5 mouse multiome data was obtained from Argelaguet et al.^43^ (GSE121708 [https://www.ncbi.nlm.nih.gov/geo/query/acc.cgi?acc=GSE121708]) and conserved rCREs were mapped onto ATAC-seq peaks in the multiome dataset using GenomicRanges findOverlaps() function. After omitting extraembryonic clusters, quadratic programming was used to order cells along the AP axis. Smoothened gene expression profiles along the AP axis were computed using TradeSeq’s predictSmooth() function.

## Statistics & Reproducibility

Statistical analyses were performed using R (version 4.3.0). Statistical details for individual experiments have been provided in the figure legends, and in methods details. No statistical methods were used to predetermine sample size. In general, three biological replicates were used for genome-wide sequencing experiments.

## Data availability

All datasets generated in this article have been deposited to the Gene Expression Omnibus under the accession code GSE224724 [https://www.ncbi.nlm.nih.gov/geo/query/acc.cgi?acc=GSE224724]. The mouse multiome data was obtained from Argelaguet et al.^43^ (GSE121708 [https://www.ncbi.nlm.nih.gov/geo/query/acc.cgi?acc=GSE121708]). Source data are provided with this paper.

## Code availability

The code used for data cleaning and analysis, along with all relevant scripts, is openly available at https://github.com/mmrothstein/tomoSeq_2025.

## Acknowledgments

The authors thank Peter Schweitzer from Cornell University BRC Genomics Core Facility for all help generating the multiome libraries and next-generation sequencing. We also acknowledge the Nikon Imaging Center at Harvard Medical School for microscopy and image analysis resources. This project was supported by National Institutes of Health grant DP2 HD102043.

## Author Contributions Statement

Conceptualization: MSC, APA, MR; Methodology: MSC, APA, MR, TK; Investigation MSC, APA, MR, TK; Visualization: MSC, MR, APA, TK; Funding acquisition: MSC; Project administration: MSC; Supervision: MSC; Writing: MSC, MR.

