## Supplementary Information for "Spatial patterning of the epigenome during vertebrate gastrulation"

**The PDF file includes:**

Supplementary Figures 1 to 11

**Other Supplementary Materials for this manuscript include the following:**

Supplementary Data 1 to 4

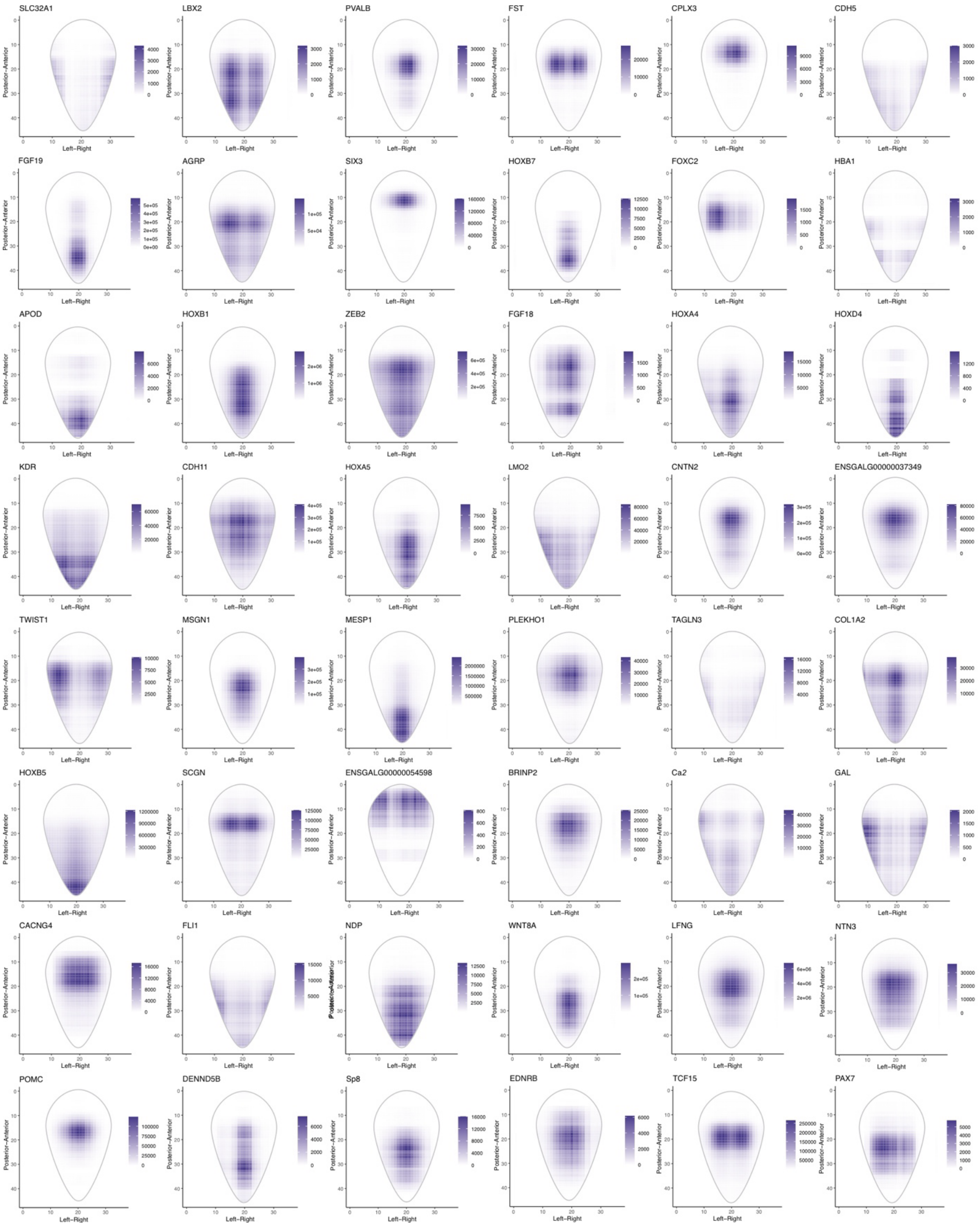
SLC32A1
LBX2
PVALB
FST
CPLX3
CDH5
FGF19
AGRP
SIX3
HOXB7
FOXC2
HBA1
APOD
HOXB1
ZEB2
FGF18
HOXA4
HOXD4
KDR
CDH11
HOXA5
LMO2
CNTN2
ENSGALG00000037349
TWIST1
MSGN1
MESP1
PLEKHO1
TAGLN3
COL1A2
HOXB5
SCGN
ENSGALG00000054598
BRINP2
Ca2
GAL
CACNG4
FLI1
NDP
WNT8A
LFNG
NTN3
POMC
DENND5B
Sp8
EDNRB
TCF15
PAX7
Posterior−Anterior
Left−Right

**Supplementary Fig. 1. 2D *in silico* prediction of gene expression patterns in the avian gastrula.** Selection of 2D projections of gene expression patterns from integration of medial-lateral and anterior-posterior tomo-Seq datasets.

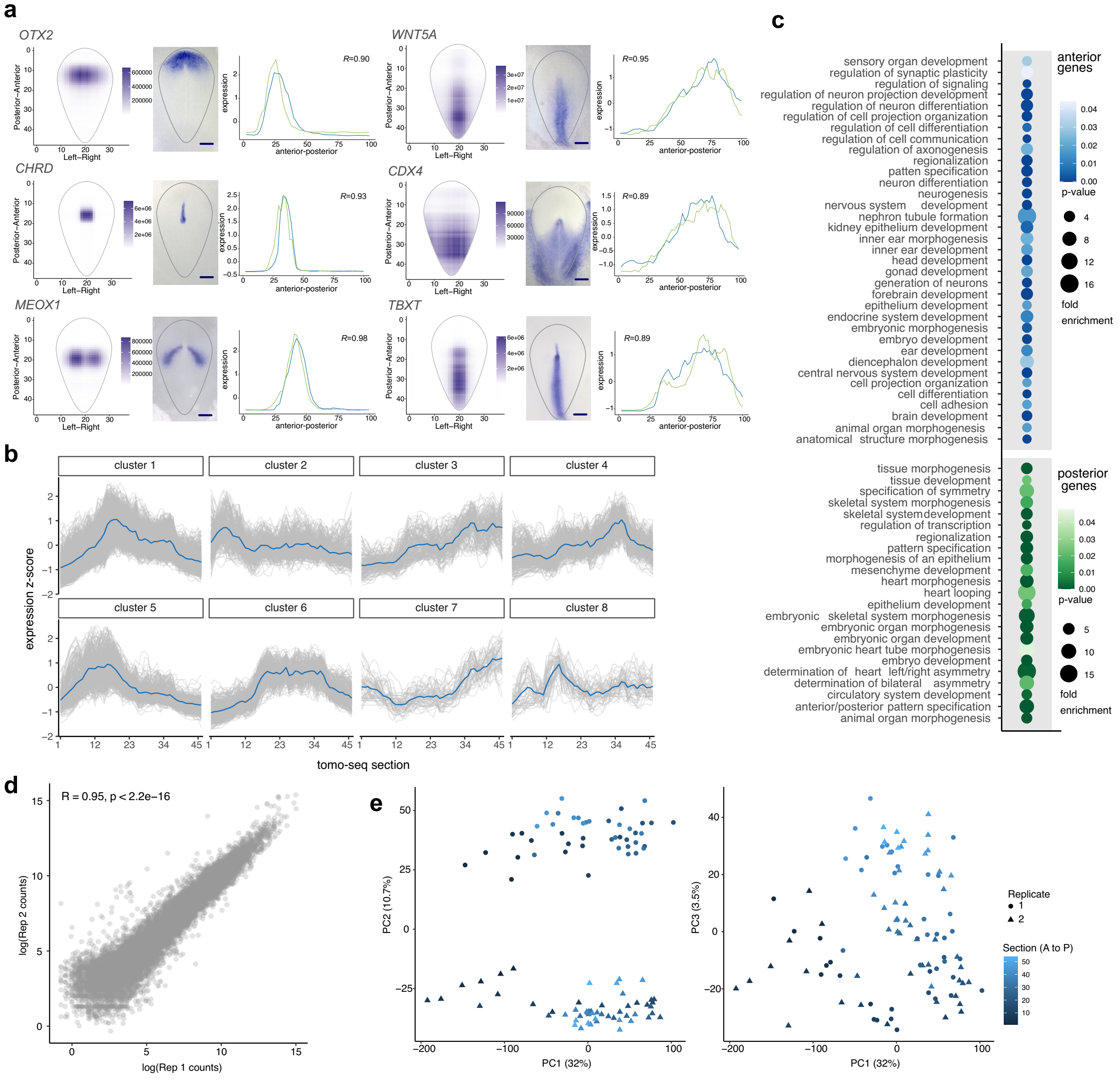


**Supplementary Fig. 2. Reconstruction of gene expression profiles via tomography-seq.** (a) *In silico* reconstruction (left) of the *OTX2*, *CHRD*, *MEOX1*, *WNT5A*, *CDX4*, and *TBXT* 2D gene expression patterns, along with the corresponding *in situ* hybridization images (center) and anterior-posterior expression profiles (right). *In silico* projections recapitulate the gene expression profiles as seen via *in situ* hybridization. Anterior-posterior expression profiles are highly correlated across biological replicates. Scale bar: 400μm. (b) Hierarchical clustering of transcriptomic data reveals dynamic expression profiles along the anterior-posterior axis. (c) Gene ontology (GO) analysis of anterior-/posterior-enriched gene sets (706 and 294 genes, respectively). GO terms associated with neurogenesis, head development, and sensory structure morphogenesis are enriched in the anterior gene set, while GO terms associated with posterior-enriched genes included mesenchymal tissue formation, including heart and skeletal morphogenesis. (d) Correlation plot showing a strong positive correlation (R=0.95) in gene expression values between tomo-Seq biological replicates. (e) Principal component analysis of tomo-Seq datasets. Principal components 1 and 3 show separation due to anterior-posterior position, while principal component 2 shows separation between biological replicates. Source data are provided as a Source Data file.

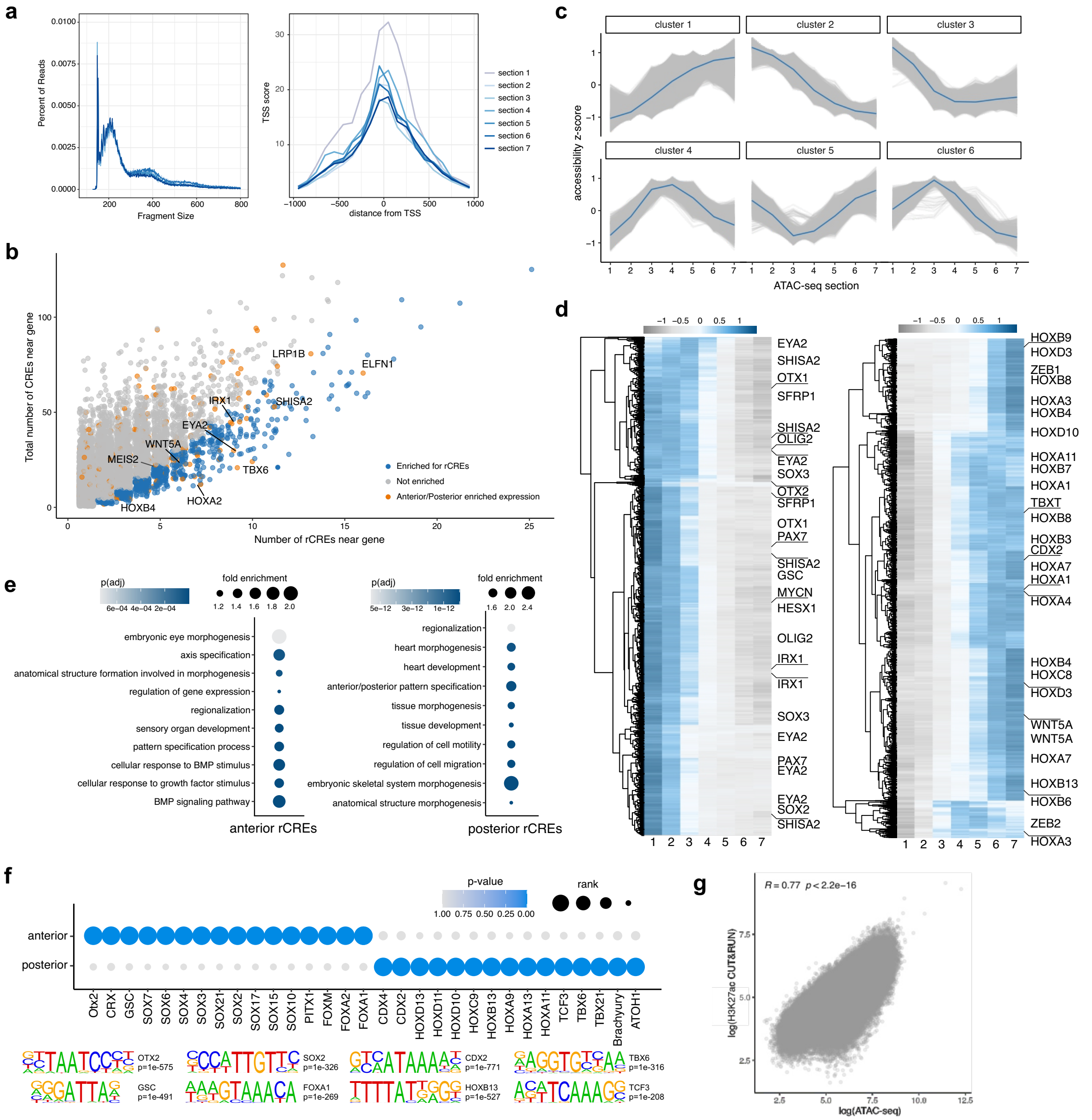


**Supplementary Fig. 3. Spatially resolved ATAC-seq in the avian gastrula.** (a) Fragment size distributions and transcription start site (TSS) enrichment plots of ATAC-seq datasets produced in 7 sections along the anterior-posterior axis. Distributions display the expected nucleosomal periodicity and enriched accessibility at annotated TSSs. (b) Enrichment plot of annotated genes showing the number of rCREs in the loci of each gene (x axis), as well as the total number of CREs in the locus of each gene (y axis). For each gene, a binomial test was conducted to determine if it is enriched for rCREs (highlighted in blue). Genes that display anterior/posterior enriched expression (as in Fig. 1f) are highlighted in orange. There is a significant association between genes enriched for rCREs in their loci and those with anterior/posterior enriched expression patterns (*P*=3.635e-06, chi-square test). (c) Hierarchical clustering of ATAC-seq datasets displays dynamic accessibility profiles along the AP axis. (d) Clustered heatmaps of significantly anterior- and posterior-enriched accessible regions (10913 and 12532 regions, respectively) as determined via TradeSeq in 7 sections along the AP axis. (e) GREAT GO analysis of anterior and posterior rCREs. Dotplots are colored by adjusted p-value and dot size represents the fold enrichment of each GO term. (f) Dotplot of top enriched transcription factor binding motifs in anterior and posterior rCREs. Anterior rCREs were enriched for motifs such as OTX2, GSC, SOX2, and FOXA1, while posterior rCREs were enriched for CDX2, HOXB13, TBX6, and TCF3 motifs. (g) Correlation plot showing a strong positive correlation (R=0.77) between ATAC-seq and H3K27AC CUT&RUN counts across all accessible peaks. Source data are provided as a Source Data file.

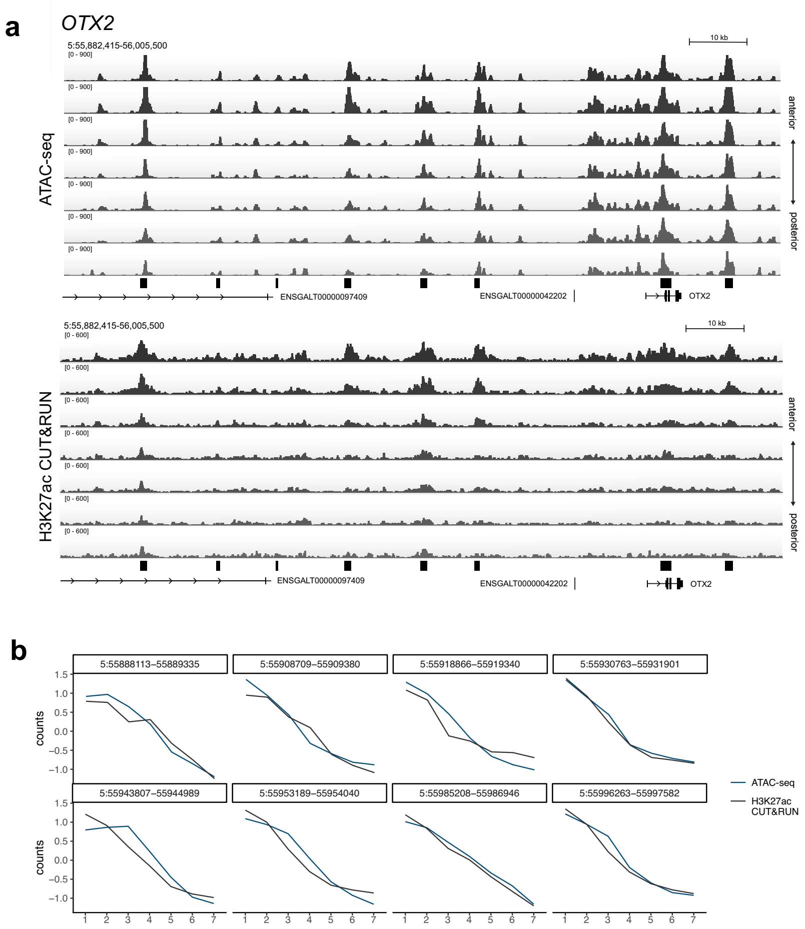


**Supplementary Fig. 4. Anterior gradients of chromatin accessibility and activity in the *OTX2* locus**. (a) Profiles of chromatin accessibility and H3K27AC CUT&RUN signal in the locus of the anterior gene, *OTX2*. (b) Quantification of chromatin accessibility and H3K27AC CUT&RUN signal at individual peaks in the *OTX2* locus. Chromatin accessibility and H3K27AC activity are highly correlated and are present in an anterior to posterior gradient. Source data are provided as a Source Data file.

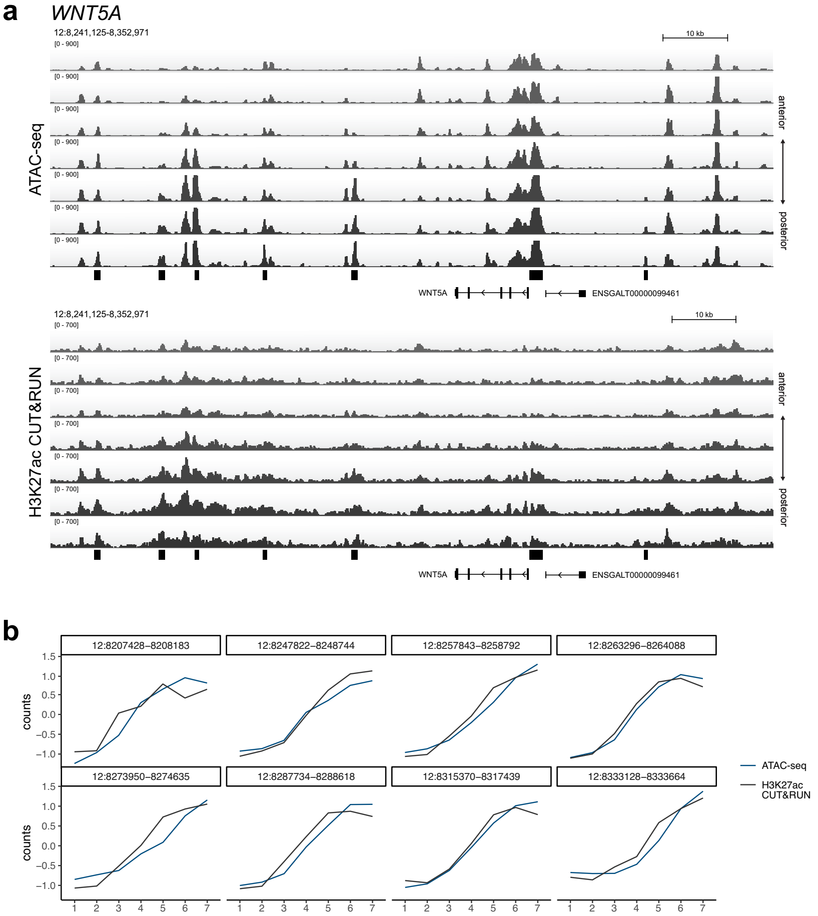


**Supplementary Fig. 5. Posterior gradients of chromatin accessibility and activity in the *WNT5A* locus**. (a) Profiles of chromatin accessibility and H3K27AC CUT&RUN signal in the locus of the posterior gene, *WNT5A*. (b) Quantification of chromatin accessibility and H3K27AC CUT&RUN signal at individual peaks in the *WNT5A* locus. Chromatin accessibility and H3K27AC activity are highly correlated and are present in a posterior to anterior gradient. Source data are provided as a Source Data file.

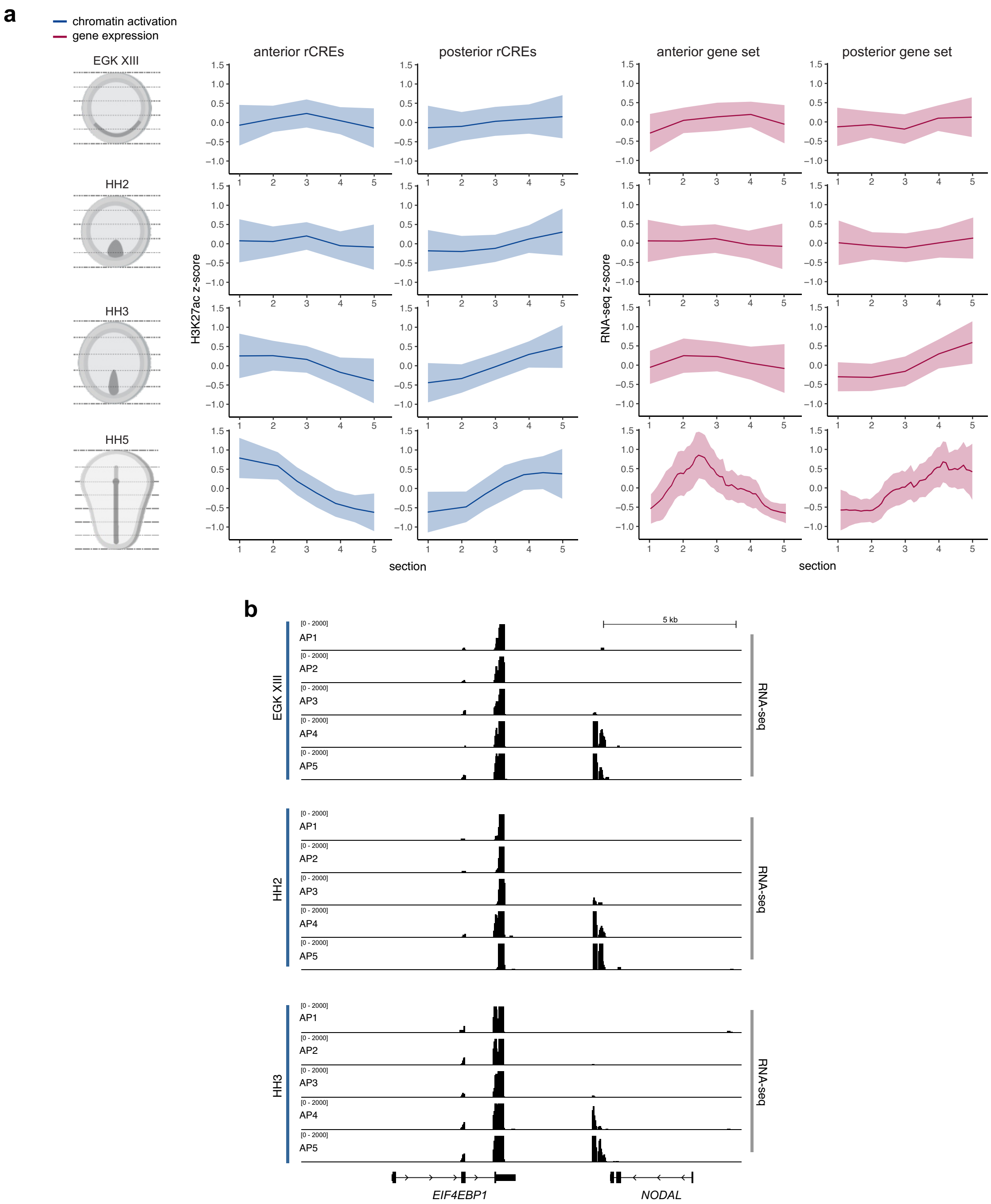


**Supplementary Fig. 6. Establishment of rCRE gradients during embryogenesis.** (a) Quantification of H3K27AC CUT&RUN signal in anterior and posterior rCREs (10913 and 12532 regions, respectively), as well gene expression values in anterior and posterior-enriched genes (706 and 294 genes, respectively) in sections along the anterior-posterior axis. Sections were collected from the early blastula (EGK XIII), late blastula (HH2), early gastrula (HH3), and late gastrula/early neurula (HH5). H3K27AC activity and gene expression are highly correlated. (b) Gene expression profiles at the *NODAL* gene locus from RNA-seq data collected in 5 sections along the AP axis at three developmental stages (EGK XIII, HH2, and HH3). RNA-seq profiles display posterior-enriched *NODAL* expression at all three timepoints. Source data are provided as a Source Data file. HH: Hamburger Hamilton.

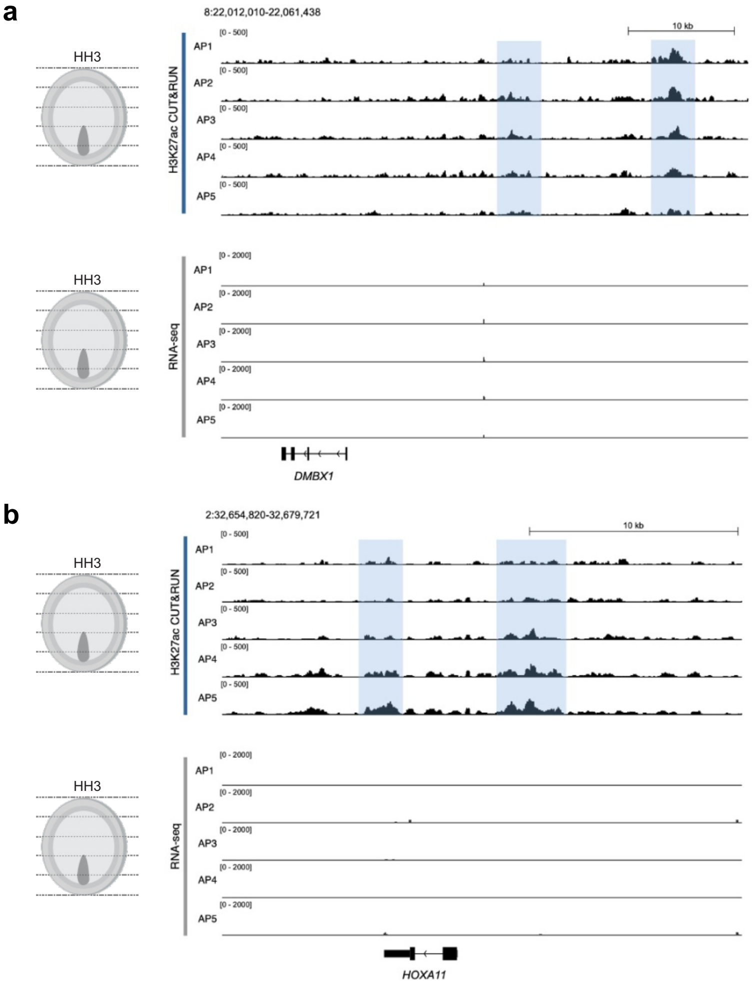


**Supplementary Fig. 7. Chromatin patterning precedes gene expression in the avian gastrula.** H3K27AC CUT&RUN and RNA-seq signal in transverse sections of HH3 avian embryos. Differential deposition of H3K27AC along the AP axis is detected before any signal mRNA signal in the *loci* of both anterior (a) and posterior (b) genes. HH: Hamburger Hamilton.

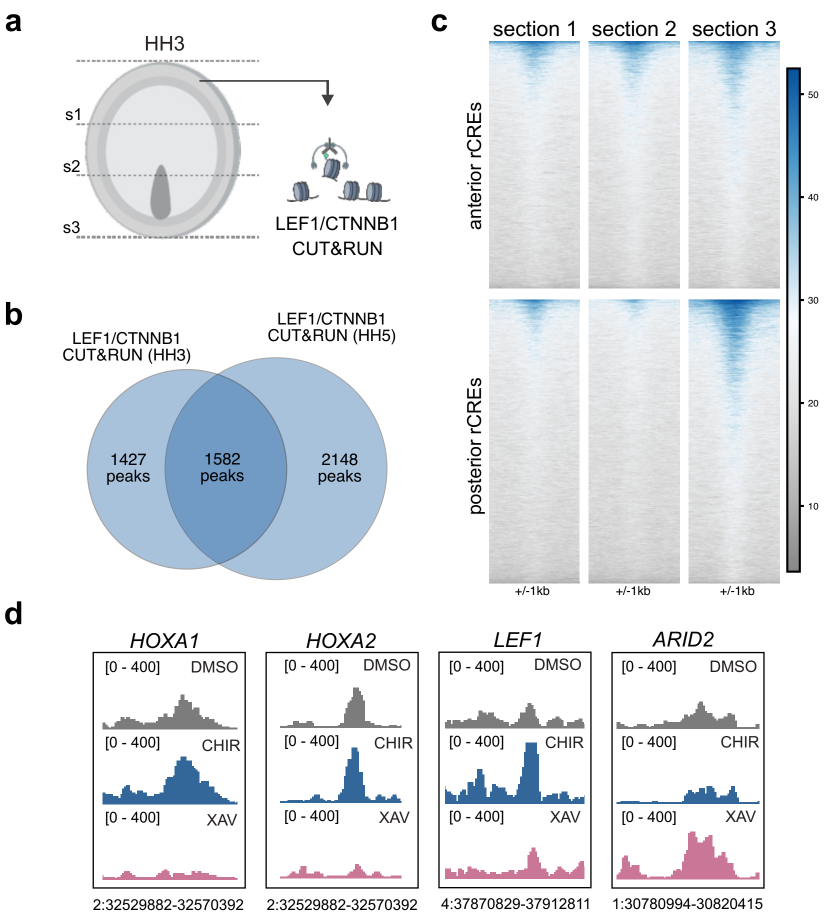


**Supplementary Fig. 8. LEF1/CTNNB1 CUT&RUN in the early gastrula**. (a) Diagram of CUT&RUN strategy targeting LEF1/CTNNB1 in anterior, medial, and posterior regions of the HH3 gastrula (n=3 per region). (b) Venn diagram displaying overlap of LEF1/CTNNB1 peaks called in HH3 and HH5 CUT&RUN datasets. (c) Tornado plots displaying LEF1/CTNNB1 signal at anterior and posterior rCREs in three serial sections of the HH3 embryo. (d) Examples of rCREs that are affected by manipulation of WNT signaling. Peaks in the *HOXA1*, *HOXA2*, and *LEF1* loci are positively regulated by WNT signaling, while a peak in the *ARID2* locus is negatively regulated by WNT signaling.

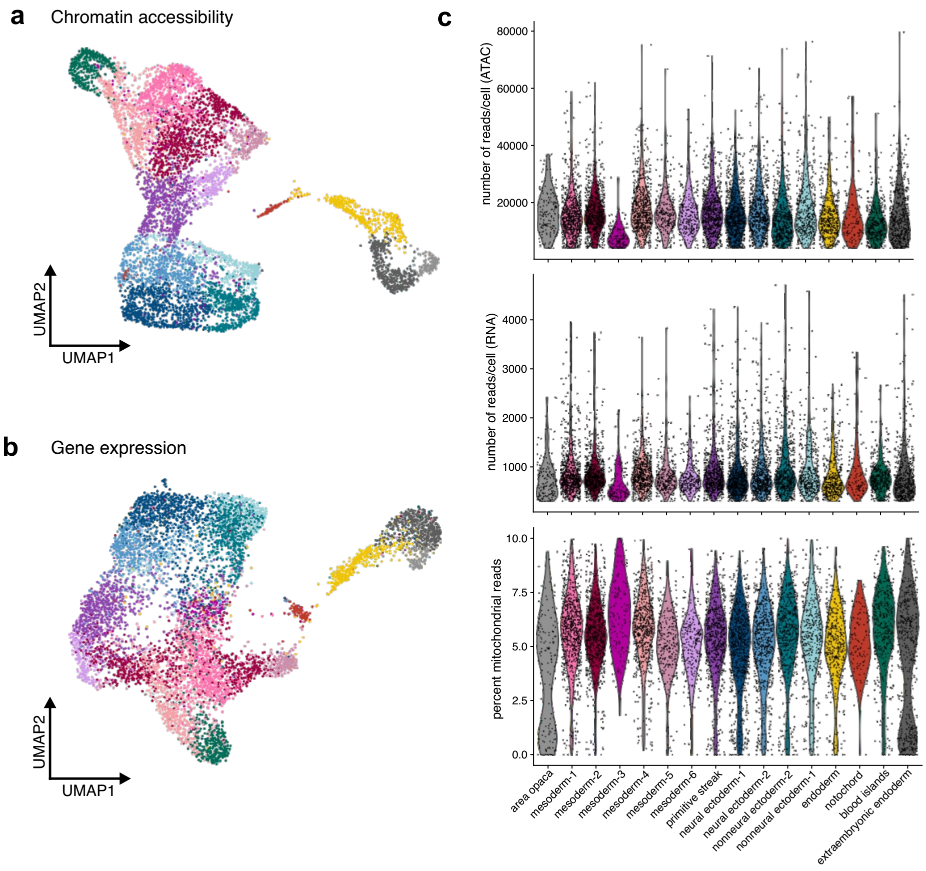


**Supplementary Fig. 9. Multiomic analysis of the avian gastrula.** (a) UMAP projection of the single-cell multiome dataset with clustering based on either chromatin accessibility or (b) gene expression. (c) Violin plots displaying quality control metrics of the multiome dataset, including the distribution of ATAC-seq and RNA-seq reads per cell, as well as the percentage of reads mapping to mitochondrial genes.

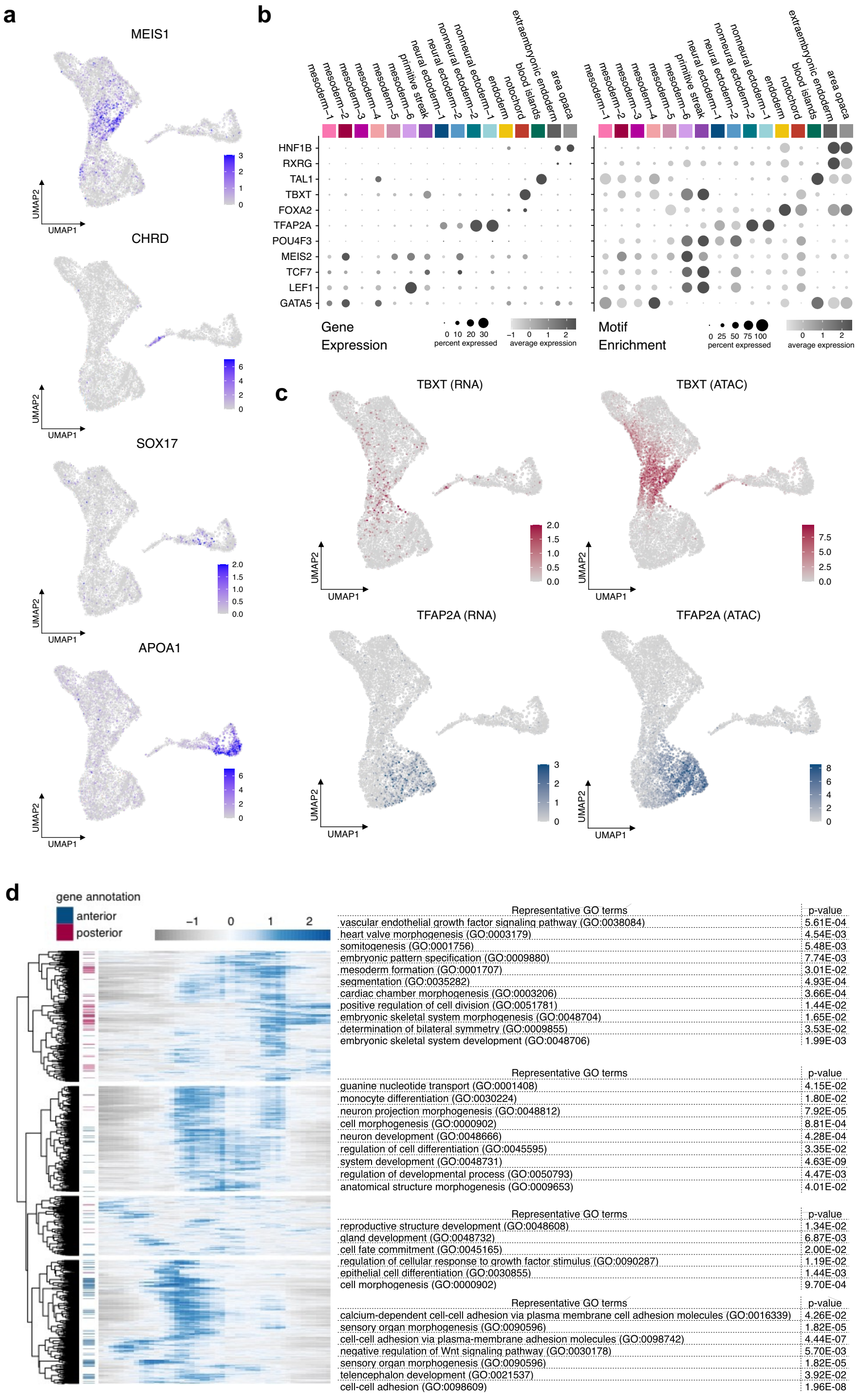


**Supplementary Fig. 10. Cluster-enriched marker genes in the single-cell multiome dataset.** (a) UMAP projections displaying gene expression enrichment via RNA-seq of tissue-specific marker genes, including the mesoderm marker *MEIS1*, the notochord marker *CHRD*, the endoderm marker *SOX17*, and the extraembryonic marker *APOA1*. (b) Dotplot displaying gene expression and motif enrichment of cluster-enriched transcription factors across single-cell multiome clusters. (c) UMAP projections displaying paired gene expression and motif enrichment values for the non-neural ectoderm marker, *TFAP2A*, and the primitive streak marker, *TBXT*. (d) Heatmap displaying anterior-posterior expression patterns of genes associated with rCREs via peak-gene linkage analysis of the HH5 multiome dataset. Anterior/posterior-enriched genes (See Figure 1f) are indicated in blue and magenta, respectively. Gene ontology (GO) analysis of four gene modules based on k-means clustering is shown with representative GO terms for each cluster. Full GO results are provided in the source data file.

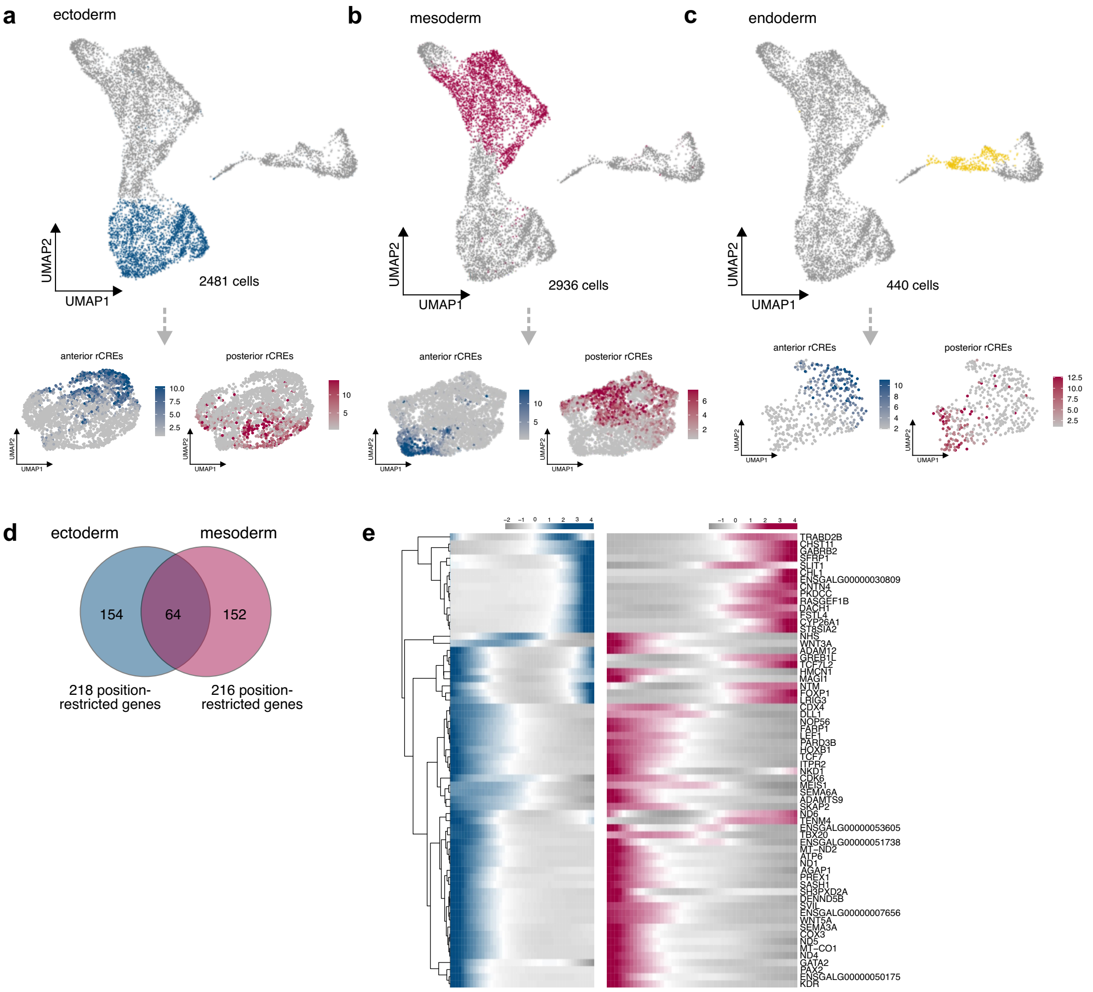


**Supplementary Fig. 11. Enrichment of rCREs in individual germ layers of the avian gastrula.** (a-c) Subsetting of the single-cell multiome dataset into three germ layer-specific subsets, including the ectoderm (2481 cells), mesoderm (2936 cells), and endoderm (440 cells) subclusters. Projection of anterior and posterior-enriched rCREs (10913 and 12532 regions, respectively) onto germ layer-specific subclusters reveals distinct subpopulations enriched for axial-restricted peaks. (d) Venn diagram displaying overlap in genes significantly associated with AP-restricted expression in the ectoderm and mesoderm subclusters. 64 genes are AP-restricted in both cell types. (e) Clustered heatmaps showing the expression of 64 genes significantly associated with AP-restricted expression in both ectoderm and mesoderm along the AP axis.
